# An activity-based labelling method for the detection of ammonia and methane-oxidizing bacteria

**DOI:** 10.1101/2021.01.14.426632

**Authors:** Dimitra Sakoula, Garrett J. Smith, Jeroen Frank, Rob J. Mesman, Linnea F.M. Kop, Mike S.M. Jetten, Maartje A.H.J. van Kessel, Sebastian Lücker

## Abstract

The advance of metagenomics in combination with intricate cultivation approaches has facilitated the discovery of novel ammonia- and methane-oxidizing microorganisms, indicating that our understanding of the microbial biodiversity within the biogeochemical nitrogen and carbon cycles still is incomplete. However, the *in situ* detection and phylogenetic identification of novel ammonia- and methane-oxidizing bacteria remains a challenge. Here, we describe an activity-based protein profiling protocol allowing cultivation-independent unveiling of ammonia- and methane-oxidizing bacteria. In this protocol, 1,7-octadiyne is used as a bifunctional enzyme probe that, in combination with a highly specific alkyne-azide cycloaddition reaction, enables the fluorescent or biotin labelling of cells harboring active ammonia and methane monooxygenases. The biotinylation of these enzymes in combination with immunogold labelling reveals the subcellular localization of the tagged proteins, while the fluorescent labelling of cells harboring active ammonia or methane monooxygenases provides a direct link of these functional lifestyles to phylogenetic identification when combined with fluorescence *in situ* hybridization. Furthermore, we show that this activity-based labelling protocol can be successfully coupled with fluorescence-activated cell sorting for the enrichment of nitrifiers and methanotrophs from complex environmental samples, facilitating the retrieval of their high quality metagenome-assembled genomes. In conclusion, this study demonstrates a novel, functional tagging technique for the reliable detection, identification, and enrichment of ammonia- and methane-oxidizing bacteria present in complex microbial communities.

## Introduction

Over the last two decades, metagenomic approaches resulted in the identification of novel groups of ammonia- (1-3) and methane-oxidizing microorganisms (4-6), highlighting that our understanding of the microbial biodiversity within the nitrogen and carbon biogeochemical cycles can still be expanded (7). However, even when detected in metagenomic datasets, linking a function to a specific microorganism remains challenging, as it often requires tedious and intricate cultivation techniques to isolate these often slow-growing and fastidious microorganisms. Thus, there is an urgent need for robust cultivation-independent methods that provide reliable information regarding the identity and activity of the microorganisms present in complex microbial communities. To achieve this, various *in vivo* and *in vitro* activity-based protein profiling (ABPP) protocols have been developed to link specific functions to catalytically active proteins (8).

ABPP techniques employ bifunctional enzyme probes that feature (i) a reactive group, which binds to the active site and thereby inhibits and covalently labels the enzyme, and (ii) an ethynyl or azide group, which can subsequently be used for the attachment of a reporter group (e.g., fluorophores or biotin) to the enzyme via a Cu(I)-catalyzed alkyne-azide cycloaddition (CuAAC) reaction (9). The use of rapid CuAAC reactions performed under mild, aqueous conditions guarantees minimal unspecific reactivity of the reporters and enables the use of low molecular weight bifunctional probes that can easily penetrate biological membranes (10-12). Subsequently, depending on the reporter type, the reporter-conjugated enzyme can be subjected in numerous downstream applications such as fluorescent imaging (13-15), mass spectrometry-based proteomics (16, 17) and affinity purification of the labelled proteins (18). Several microbial proteins have been studied using ABPP protocols so far. These studies provided significant insights into antibiotic resistance, enzymatic functions in pathogenic bacteria (e.g., serine proteases, kinases, ATPases, fatty acid synthases, glycoside hydrolases) and protein redox dynamics (8). However, only a limited number of ABPP protocols targeting microbial proteins catalyzing key processes of biogeochemical cycles have been developed (2, 19-21).

Autotrophic ammonia- and methane-oxidizing bacteria (AOB and MOB, respectively) (22) are ubiquitous in the environment (4) and are of high biotechnological interest (23-29). Besides their similar environmental niches, they share many biochemical, morphological and physiological characteristics (30). More specifically, both microbial guilds can perform aerobic oxidation of ammonia and methane due to the substrate promiscuity of their key enzymes, but neither group exhibits growth on the alternative substrate (30-32).

The ammonia monooxygenase (AMO) belongs to the protein family of copper-containing membrane-bound monooxygenases and is responsible for the first oxidation step of ammonia to hydroxylamine in AOB and ammonia-oxidizing archaea (AOA). Similarly, the two different forms of methane monooxygenases (MMOs) identified in MOB can oxidize methane to methanol (33, 34). The membrane-bound particulate MMO (pMMO) exhibits high genetic, structural and catalytic similarities to AMO enzymes (35-37) and is present in most MOB. However, some MOB also contain a cytoplasmic iron-containing soluble MMO (sMMO), which is expressed mainly under copper-limited conditions (38, 39) and can even constitute the sole MMO in some MOB (40). Although the substrate for both MMOs is identical, the sMMO shares little structural homology with the pMMO and AMO enzymes (37). In recent years, some advances have been made in studying these enzymes (33, 41). However, the recent discovery of the novel comammox (<u>com</u>plete <u>amm</u>onia <u>ox</u>idation) *Nitrospira*, featuring a distinct AMO enzyme previously misclassified as an “unusual” pMMO (1, 2), exemplified our still incomplete understanding of the microbial diversity catalyzing biological ammonia and methane oxidation.

To date, many insights into the distribution, abundance and activity of ammonia-oxidizing prokaryotes (AOP) and MOB in natural environments have been obtained by employing reversible (42-45) and irreversible inhibitors (19, 46-48). Many terminal *n-*alkynes can be catalytically activated by AMO and MMOs, producing reactive intermediates that bind covalently to the enzyme and act as mechanism-based irreversible inactivators (47, 49-52). The alkyne 1,7-octadiyne (1,7OD) was recently characterized as a mechanism-based inactivator of the AMO enzyme in *Nitrosomonas europaea* and was successfully used in *in vitro* ABPP protocols (19).

In this study, we describe the use of 1,7OD in an ABPP-based protocol for the *in situ* biotin or fluorescent labelling of bacterial AMO and MMOs. In combination with transmission electron microscopy, the biotin tagging of the enzymes permits the study of their intracellular distribution, while their fluorescent tagging allows the function-based detection of catalytically active ammonia and methane-oxidizing microorganisms by fluorescence microscopy and can be combined with fluorescence *in situ* hybridization (FISH) for phylogenetic identification. Moreover, this method can be efficiently combined with fluorescence-activated cell sorting for the enrichment of ammonia- and methane-oxidizing bacteria from complex environmental samples. Combined with downstream metagenomics, this facilitates the targeted retrieval of high quality metagenome-assembled genomes (MAGs) of a functionally-defined subpopulation.

## Materials and methods

### Cultivation

A pure culture of the canonical ammonia-oxidizing bacterium *Nitrosomonas europaea* (ATCC 25978) was obtained from the Leibniz Institute DSMZ-German Collection of Microorganisms and Cell Cultures (Braunschweig, Germany). *N. europaea* was grown in batch in mineral salts medium amended with 10 mM NH_4_Cl (53). Cultures of the methane-oxidizing bacteria *Methylotetracoccus oryzae* (nitrogen-fixing, Type Ib methanotroph), *Methylosinus sporium* M29 (Type II methanotroph), *Methylacidiphilum fumariolicum* (Type III methanotroph) and *Methylocella tundrae* (Type II methanotroph containing sMMO only) were maintained in batch in mineral salts medium as previously described (6, 54-56). A pure culture of the canonical nitrite-oxidizing bacterium *Nitrospira moscoviensis* M-1 (DSM 10035) was grown in a mineral salts medium according to Mundiger *et al*. (57). *Nitrospira inopinata* cells were grown as described elsewhere in the presence of 1mM NH_4_Cl (58). A pure culture of the diazotrophic *Kyrpidia spormannii* FAVT5 was cultivated according to Hogendoorn *et al*. (59). An enrichment culture containing the comammox species *Ca*. Nitrospira nitrosa and *Ca*. Nitrospira nitrificans was maintained in a sequencing batch bioreactor as reported previously (2). An enrichment culture of the novel comammox species *Ca*. Nitrospira kreftii was maintained in a membrane chemostat as described elsewhere (60). A nitrifying enrichment culture containing canonical AOB, canonical (i.e., lacking AMO) and comammox *Nitrospira*, as well as non-nitrifying microorganisms was maintained in a continuous membrane bioreactor in mineral salts medium under substrate-limited conditions. Activated sludge samples were obtained from the aeration tank of a municipal wastewater treatment plant (wwtp) in Groesbeek, The Netherlands (51°45’34.3”N 5°57’15.9”E).

Unless stated otherwise, biomass was harvested from the bioreactors and wwtp by gentle centrifugation (600 × *g* for 10 min), washed twice, and subsequently resuspended in mineral salts medium to a final density corresponding to approximately 100 μg protein ml^-1^.

### Inactivation of comammox Nitrospira by 1,7OD

To test for the irreversible inactivation of comammox *Nitrospira* by 1,7OD, biomass from the *Ca*. N. kreftii enrichment culture (60) was harvested and resuspended in 50 ml mineral salts medium without nitrogen source containing 100 μΜ 1,7OD (99% purity, Merck KGaA, Darmstadt, Germany) from a stock solution (75.4 mM) in dimethyl sulfoxide (DMSO). Cells were incubated for 10 minutes in the dark in a shaking incubator (150 rpm, 24°C). Subsequently, incubations were supplemented with ammonium to a final concentration of 350 μM and were incubated for 72 hours. Control incubations receiving only 66.4 μl DMSO without 1,7OD were performed in parallel. As a positive control for the inhibition of the ammonia oxidation activity, cells were incubated in the presence of 100 μM allylthiourea (ATU; ≥98% purity, Merck KGaA, Darmstadt, Germany), which has been shown to inhibit comammox *Nitrospira* previously (2). Finally, the specificity of 1,7OD as inhibitor of the ammonia-oxidizing activity was verified by incubating comammox *Nitrospira* cells in the presence of 100 μM 1,7OD and 20 μM nitrite. All incubations were performed in three biological replicates. During incubation, samples for the determination of ammonium, nitrite and nitrate concentrations were taken every two hours for the first 8 hours of incubation, as well as after 24, 48 and 72 hours of incubation.

### Analytical methods

Ammonium concentrations were measured fluorometrically using a modified orthophatal-dialdehyde assay (61). Nitrite and nitrate concentrations were determined colorimetrically with the Griess reaction (62) as described elsewhere (63). For the determination of total protein content, cells were lysed using the Bacterial Protein Extraction Reagent (B-PER™, Thermo Fisher Scientific, Waltham, MA, USA) according to manufacturer’s instructions in combination with mild sonication (1 min, 20 Hz). Post lysis, protein concentrations were measured using the Pierce bicinchoninic acid protein assay kit (Thermo Fisher Scientific, Waltham, MA, USA) according to the ‘Enhanced Test-tube’ protocol. All fluoro- and colorimetric measurements were performed using a Tecan Spark^®^ M10 plate reader (Tecan Trading AG, Männedorf, Switzerland).

### In situ ABPP-based fluorescent labelling of AMO and MMO

Active biomass was harvested as described above, resuspended in 50 ml mineral salts medium and incubated with 100 μM 1,7OD for 30 minutes (150 rpm) in the dark. Subsequently, cells were pelleted by gentle centrifugation (600 × *g* for 10 minutes), washed twice in sterile PBS, pH 7.5 and fixed using a 50% (v/v) ethanol/PBS solution for 10 minutes at room temperature (RT). Fixed biomass was washed once with PBS and subjected to the CuAAC reaction, which was performed in plastic microcentrifuge tubes (1,5 ml) in a final volume of 250 μl. Biomass was resuspended in 221 μl sterile PBS, mixed with 12.5 μl of a 100 mM freshly prepared sodium ascorbate solution (≥99% purity, Merck KGaA, Darmstadt, Germany) and 12.5 μl of a 100 mM freshly prepared aminoguanidine hydrochloride solution (≥98% purity, Merck KGaA). A dye mixture containing 1.25 μl of a 20 mM CuSO_4_ solution (99,99% purity, Merck KGaA), 1.25 μl of a 100 mM Tris(3-hydroxypropyltriazolylmethyl)amine solution (THPTA; 95% purity, Merck KGaA) and 0.3 μl of 5 mM Azide-Fluor 488 in DMSO (≥90% purity, Merck KGaA) was incubated in the dark for 3 minutes. Subsequently, the dye mixture (2.8 μl) was added to the CuAAC reaction tubes. The tubes were gently mixed and incubated for 60 minutes (RT, in the dark). CuAAC reactions were terminated by harvesting the cells by gentle centrifugation (600 × *g* for 10 minutes). Cell pellets were washed three times with sterile PBS to remove unbound fluorophores and either used directly for downstream analyses, or resuspended in a 50% (v/v) ethanol/PBS solution and stored at -20°C.

### Fluorescence in situ hybridization and microscopy

AMO/MMO labelled biomass was hybridized with fluorescently labelled oligonucleotides as described by Daims *et al*. (64). Probes used in this study (Table S1) were 5’ or 5’ and 3’-labelled with the dyes Cy3 or Cy5 (65). After hybridization and washing, slides were dried and embedded in DAPI-containing Vectashield^®^ antifading mounting medium (Vector Laboratories Inc., Burlingame, CA). Probe and ABPP-conferred fluorescence was recorded using a Leica TCS Sp8x confocal laser microscope (CLSM; Leica Microsystems B.V., Amsterdam, the Netherlands) equipped with a 405 nm UV diode and a pulsed white light laser. Images were recorded using 63× or 100× oil immersion objectives at a resolution of 1,024 × 1,024 pixels and 8-bit depth.

The spatial distribution of the ABPP-based AMO/MMO fluorescent signal within bacterial cells was investigated using the HyVolution deconvolution module of the Huygens Essential Suite (Scientific Volume Imaging B.V, Hilversum, The Netherlands). *N. europaea, N. inopinata, M. oryzae* and *M. tundarae* were used as representatives of canonical AOB, comammox bacteria, and pMMO and sMMO-only containing MOB, respectively. Images were acquired using a Leica Sp8x CLSM with a 100× oil immersion objective, a 0.5 AU pinhole size at a resolution of 1,024 × 1,024 pixels, and were deconvolved using the resolution-optimized algorithm of the HyVolution module.

### Immunogold localization of the AMO and pMMO enzymes and quantification

Active cells of *N. europaea* and *M. oryzae* were inactivated using 1,7OD as described above, harvested, fixed for 30 minutes at RT (2% PFA, 0.5% GA in 0.1 M phosphate buffer pH 7) and processed for Tokuyasu sectioning (66, 67). Cells that were incubated in the presence of DMSO without 1,7OD were used as a control. After trimming in order to provide a suitable block face, 65 nm sections were cut on a 35° diamond knife (Diatome, Biel/Bienne, Switzerland) at -100°C in a cryo-ultramicrotome (UC7/FC7 Leica Microsystems, Vienna, Austria), picked up using a mixture of 1% methylcellulose in 1.2 M sucrose (68) and transferred to 100#H copper grids (Agar Scientific LTD., Essex, UK).

In order to successfully localize the AMO/pMMO enzymes in the bacterial cells, the thawed cryo-sections were subjected to a CuAAC reaction (as described above) using 5 mM biotin-azide in DMSO (> 95% purity, Jena Bioscience GmbH, Jena, Germany) instead of Azide-Fluor 488. Subsequently, immuno-gold localization of the biotinylated enzymes was performed according to Slot and Geuze (2007) using a rabbit anti-biotin antibody (ab53494, Abcam plc., Cambridge, UK, diluted 1:600 in 1% BSA-C) and 10 nm protein-A gold (CMC-UMC Utrecht, the Netherlands). Sections were embedded in methyl-cellulose containing 0.2% uranyl-acetate before imaging using a JEOL JEM-1400 flash transmission electron microscope, operating at 120 KV.

The localization of the immuno-gold labels was counted in nine to twelve cells for each sample and was classified in one out of four classes: i) resin, ii) membrane-associated, iii) cytoplasmic and iv) unclear. To this end, circles with a diameter of 15 nm were drawn around the center of each gold particle. The gold particles themselves had a given diameter of 10 nm and the antibody-protein A complex accounted for another 10 nm, thus the epitopes the antibodies were bound to were expected to lie somewhere within these 15 nm circles. In case gold particles were located in the cytoplasm but with a membrane present within the circle, the label localization was classified as unclear.

The location of the epitopes that classified as ‘membrane-associated’ was investigated in greater detail. More specifically, labels on the periplasmic side of the intracytoplasmic membrane compartments were classified as ‘periplasmic’ while labels present on the cytoplasmic side were classified as ‘cytoplasmic’. Labels localized on transversely sectioned membrane stacks or exactly on top of membranes, were classified as unclear.

### Effect of growth stage on ABPP-based fluorescent labelling efficiency

Cells of *N. europaea* were grown as described above in buffered medium (4.8 g/L HEPES). Control incubations with *N. europaea* cells grown in the absence of ammonium as well as abiotic controls were performed in parallel. All incubations were performed in two biological replicates. Samples (10 mL) were taken regularly to monitor protein content, ammonium concentration and pH. Additional samples (50 mL) were subjected to the ABPP-based labelling protocol as described above. For the quantification of the relative biovolume fractions that depicted an AMO labelling, as well as the intensity of the signal, the ABPP-based protocol was combined with FISH using an equimolar mixture of the probes NSO190, NSO1225 and NEU653 (Cy3) and EUB338mix (Cy5) (Table S1). Subsequently, 15 image pairs were recorded per sample at random fields of view using the Leica TCS Sp8x CLSM. The images were imported into the image analysis software *daime* (69) and evaluated as described elsewhere (70).

### Targeted cell sorting of AMO/MMO-containing bacterial cells

Biomass from a nitrifying enrichment culture and activated sludge from a municipal wwtp were subjected to the ABPP-based AMO/MMO labelling protocol. For both biomasses, three types of control incubations were performed in parallel. These contained (i) 1,7OD without the addition of the fluorescent azide-labelled dye in the CuAAC reaction, (ii) DMSO without 1,7OD during the initial incubation, with complete subsequent CuAAC reaction, and (iii) DMSO without 1,7OD during the initial incubation and without addition of the fluorescent azide-labelled dye in the CuAAC reaction. After labelling, biomass was dispersed by mild sonification (30 s at 20 Hz) and stored at -80°C in glycerol-TE (GlyTE) buffer (10 mM Tris-HCl, 1 mM EDTA pH 8.0, 5% v/v glycerol) until further processing. Samples were thawed on ice and subjected to fluorescence-activated cell sorting (BD FACSMelody^TM^, BD Biosciences, NJ, USA). Based on the initial cell density determined by FACS, samples were diluted up to 1:4 with sterile PBS to ensure a final event rate of <10.000 events sec^-1^. Control incubations were used to determine a gating strategy for the specific sorting of the labelled population in the samples that best excluded unlabeled and autofluorescent cells. Subsequently, DNA was extracted from the bulk-sorted cell clusters, the unsorted control incubations and the initial biomass, followed by metagenomic sequencing.

Additionally, we tested for potential biases on the community composition caused by treatment steps unrelated to the ABPP-based protocol. These included potential effects of washing steps, biomass fixation, sonication and storing conditions. For this purpose, biomass from the nitrifying enrichment culture was subjected to different combinations of physical and chemical treatments occurring during the ABPP-based protocol (Table S2), followed by DNA isolation and metagenomic sequencing (see below).

### DNA extraction and metagenomic sequencing

Total DNA from all samples was extracted using the DNeasy Blood & Tissue Kit (Qiagen Ltd., West Sussex, UK) according to manufacturer’s instructions. Genomic sequencing libraries were prepared using the Nextera XT Kit (Illumina, San Diego, CA, USA) following manufacturer’s recommendations using a total of 1 μg of input DNA, normalized to a concentration of 0,2 ng/µl. The libraries were sequenced using an Illumina MiSeq with MiSeq Reagent Kit v.3 (2x 300 bp, Illumina, San Diego, CA, USA) according to manufacturer’s instructions.

### Metagenome assembly and binning

Quality-trimming, adapter removal and contaminant filtering of Illumina paired-end sequencing reads was performed using BBDuk (BBTools version 37.76; with settings ktrim=r, k=23, mink=11, hdist=1, qtrim=rl, trimq=17, maq=20, maxns=0, minlen=150) (71). Processed reads from all libraries from the same biological sample were *de novo* co-assembled using metaSPAdes v3.11.1 (72) with default settings, which iteratively assembled the metagenome using *k*-mer sizes of 21, 33, 55, 77, 99 and 127 bp. After assembly, reads were mapped back to the metagenomic contigs for each sample separately using Burrows-Wheeler Aligner (BWA version 0.7.17) (73), employing the “mem” algorithm. The mapping files were processed and converted using SAMtools version 1.9 (74). Metagenomic binning was performed including all contigs ≥1,500 bp, using differential coverage, sequence composition and linkage information. To optimize the binning results, five different binning algorithms were used: BinSanity v0.2.6.1 (75), COCACOLA (76), CONCOCT (77), MaxBin 2.0 v2.2.4 (78) and MetaBAT 2 v2.12.1 (79). The five bin sets were supplied to DAS Tool v1.0 (80) for consensus binning, yielding the final optimized bins. Bin quality and completeness were assessed based on single-copy marker gene presence by CheckM v1.0.7 (81). Phylogenetic assignment of the final bins was predicted using the classification workflow of GTDB-Tk v0.2.1 (82).

Besides the assembling and binning method described above, the sequencing data retrieved from the FACS-sorted activated sludge sample were assembled separately, with subsequent binning of all contigs ≥1,500 bp and manual refinement using the anvi’o metagenomics workflow (83). Here, contigs not clustering with the rest of the bin, based on sequence composition or coverage, were inspected individually in the assembly graph using Bandage v0.8.1 (84). Additionally, genes present on these contigs were compared against the NCBI non-redundant nucleotide database using BLASTx (85). Contigs without connections in the assembly graph and containing genes not matching the taxonomic affiliation of the bin were removed from the respective bin. In order to achieve a consistent naming, these bins were linked back to the ones obtained by the automated binning approach described above using dRep (v2.0.0.) with a 99% identity cutoff (86).

For all samples, the total number of sequenced base pairs (bp) and reads, the number of aligned bp per contig, and the number of mapped reads per contig was obtained as described above. Final bins were annotated using PROKKA v1.12-beta (87) against the RefSeq bacterial non-redundant protein database (release 92).

### Marker gene recovery and read recruitment

Genes on assembled contigs ≥1,500 bp were identified by Prodigal (88). The translated amino acid sequences were mined using Hidden Markov Models (HMM) available from the Pfam database to extract the assembled genes for ammonia monooxygenase subunit A (*amoA*, PF02461) and RNA polymerase beta subunit (*rpoB*, PF04563) (89, 90). Genes with a sequence score less than 20 or an e-value greater than 1e-5 were inspected manually to remove false-positives. Recovered amino acid sequences were queried (blastp version 2.10.0+ -evalue 1e-50, -culling_limit 10, -qcov_hsp_perc 90, -max_hsps 5) against the NCBI non-redundant protein database (version 5) to infer their taxonomic affiliations, and to identify gene fragments (less than 45% length of closest reference), which were excluded from quantitative analyses (91). Nucleotide sequences of the genes and assembled contigs ≥1,500 bp were used to quantify recruited reads using BBMap (BBTools version 37.76 idfilter=0.99, pairedonly=t, ambiguous=random) (71). Read recruitment per gene and sample was normalized by gene length and sequencing depth as paired Reads per Kilobase gene length per Million reads recruited to the assembly (RPKM). R was used to normalize read abundances, calculate correlations, and for visualization (92, 93).

### Data availability

Sequencing data obtained in this study have been deposited in the National Center for Biotechnology Information (NCBI) database under BioProject accession numbers PRJNA691748 (nitrifying enrichment culture) and PRJNA691751 (activated sludge sample).

## Results

### Inhibition of ammonia oxidation in comammox by 1,7OD

The ability of 1,7OD to act as an irreversible and mechanism-based inactivator of the AMO enzyme in the canonical AOB *Nitrosomonas europaea* was previously demonstrated (19), and the pMMO of Methylococcus capsulatus (Bath) was shown to be partially inhibited by alkynes with chain lengths ≤C_7_ (94). However, comammox *Nitrospira* harbor a phylogenetically distinct AMO enzyme (1, 2, 36). In order to test the efficiency of inactivation of the comammox AMO by 1,7OD, inhibition of ammonia oxidation activity was determined in the *Ca*. N. kreftii enrichment culture, which contained comammox *Nitrospira* as the sole ammonia-oxidizing microorganism (60).

In the absence of 1,7OD, the culture oxidized 350 μM ammonium in 48 hours to nitrate without intermediate nitrite accumulation. In contrast, addition of 100 μΜ 1,7OD resulted in a rapid and complete inhibition of the ammonia-oxidizing activity, which was comparable to the inhibitory effect of 100 μM ATU (Figure 1A, B). No influence of 1,7OD or ATU on the nitrite-oxidizing activity of the culture was detected (Figure 1C, D).

**Figure 1.**
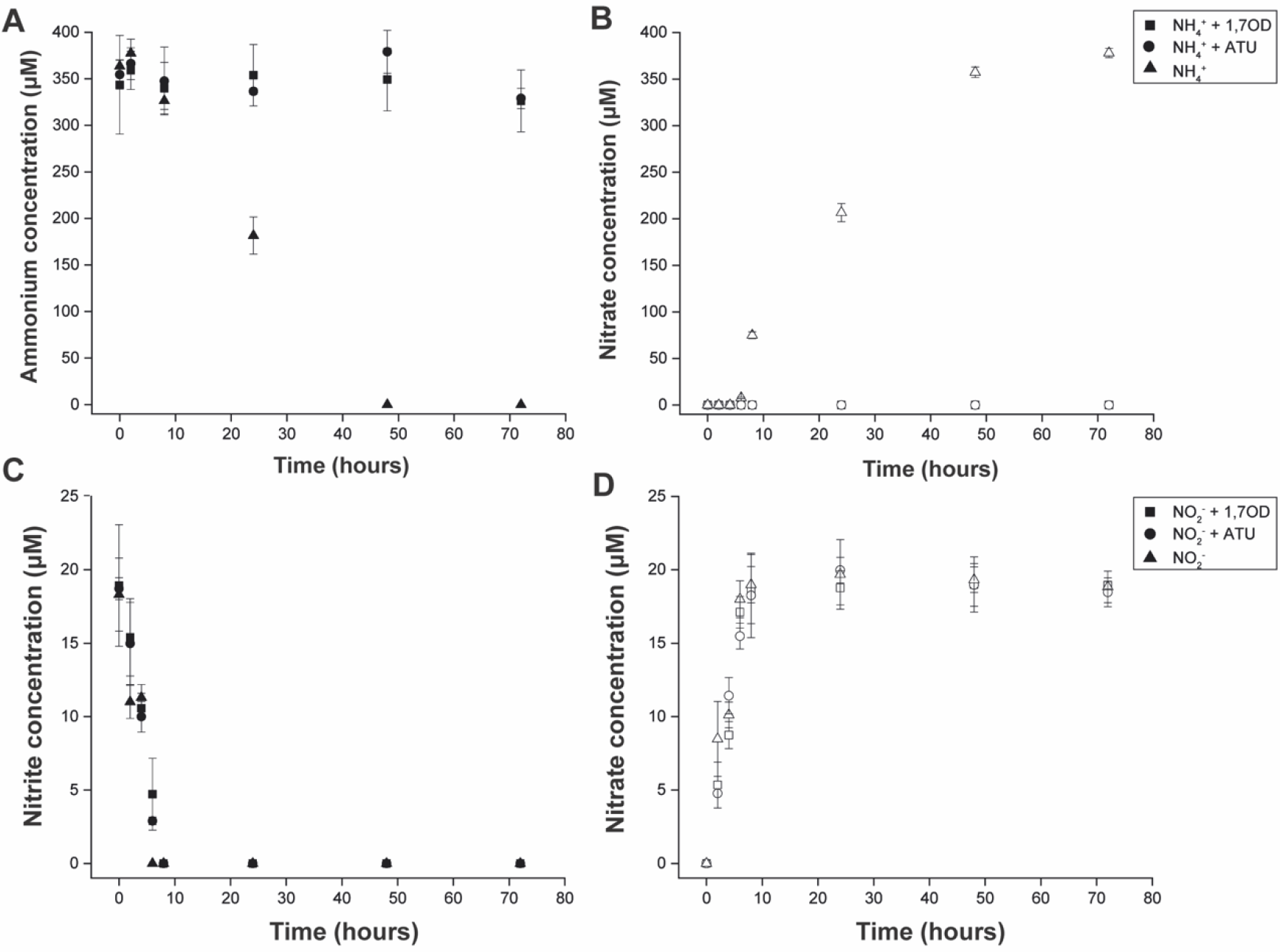
Inhibition of the ammonia oxidation activity of the comammox organism *Ca*. N. kreftii by 1,7OD. (A) Ammonium consumption and (B) nitrate production in the presence of 350 μM NH4 ^+^ with and without the addition of 100 μΜ 1,7OD or ATU. (C) Nitrite consumption and (D) nitrate production in the presence of 20 μM NO_2_ ^-^ with and without the addition of 100 μΜ 1,7OD or ATU. Concentrations of the substrates ammonium (A) and nitrite (C) are indicated by solid, produced nitrate (B, D) by open symbols. Data points represent the mean of technical triplicates, error bars the standard deviations between three biological replicates.

### In situ fluorescent labelling of AMO and MMO

We adapted the ABPP-based AMO labelling protocol developed by Benett *et al*. (19) to allow the *in situ* fluorescent labelling of ammonia and methane-oxidizing bacteria. Biomass containing different ammonia or methane oxidizers was incubated in the presence of 100 μM 1,7OD, which lead to the formation of stable 1,7OD-enzyme complexes within the bacterial cells (19, 47). After EtOH fixation, the inactivated biomass was subjected to a highly specific CuAAC reaction. The use of the CuAAC reaction enabled the covalent coupling of the alkyne-labelled enzymes to an azide-labelled Fluor488 dye, which resulted in the fluorescent labelling of AMO (Figure 2) and MMO-containing microorganisms (Figure 3).

**Figure 2.**
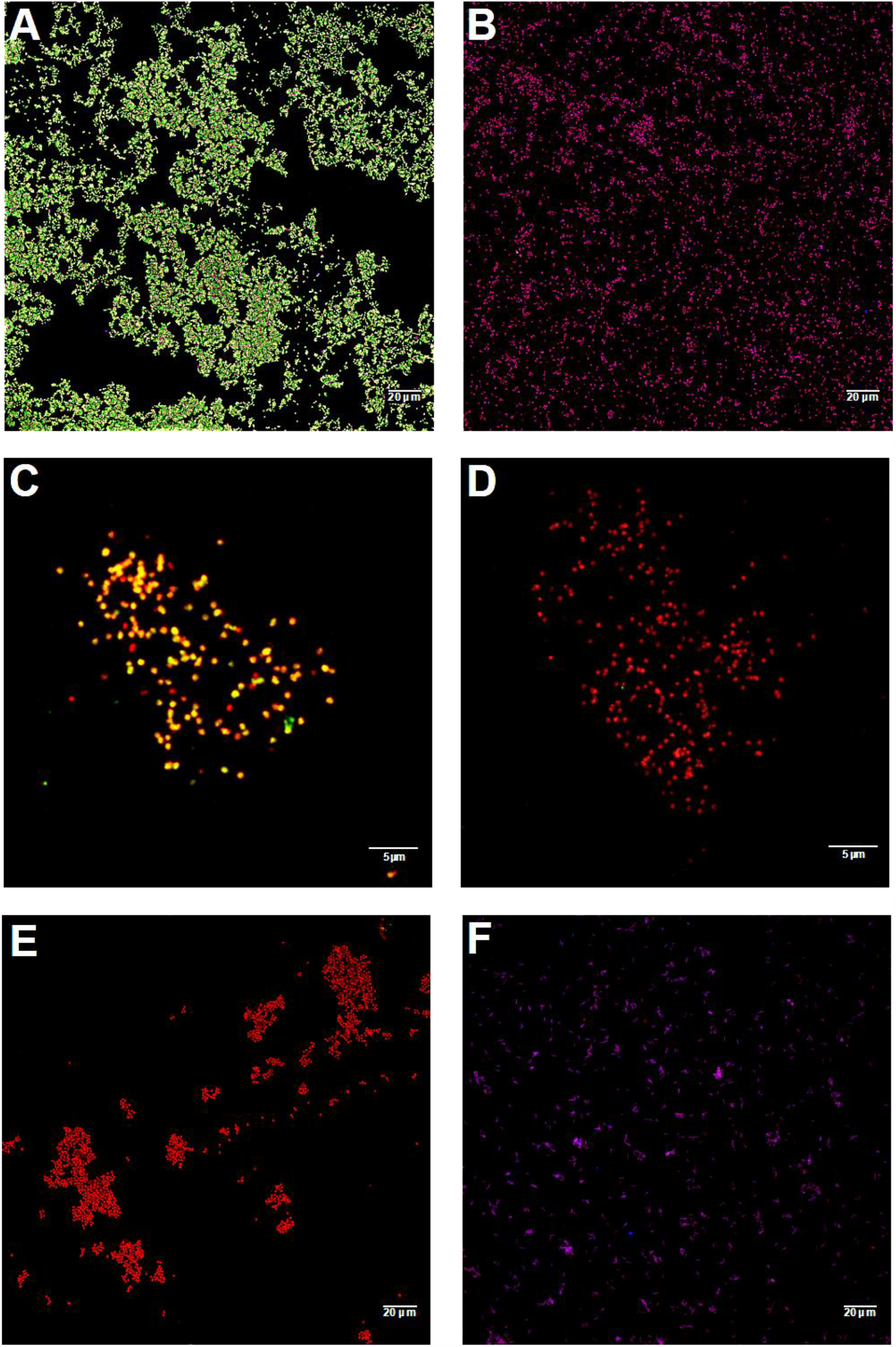
ABPP-based fluorescent labelling of catalytically active ammonia-oxidizing prokaryotes. (A, C, E, F) 1,7OD pre-incubated and (B, D) 1,7OD untreated (control) cells of (A, B) *Nitrosomonas europaea*, (C, D) *Nitrospira inopinata*, (E) *Nitrosocosmicus franklandus*, (F) *Nitrospira moscoviensis*. Cells are stained with the AMO/MMO-labelling protocol (green) and FISH probes specific for (A, B) AOB (Nso190, Neuo653, Nso1225; red), (C, D, F) *Nitrospira* (Ntspa662, Ntspa712; red), (E) AOA (Arch915; red) and (A, B, F) all bacteria (EUB338mix; blue).

**Figure 3.**
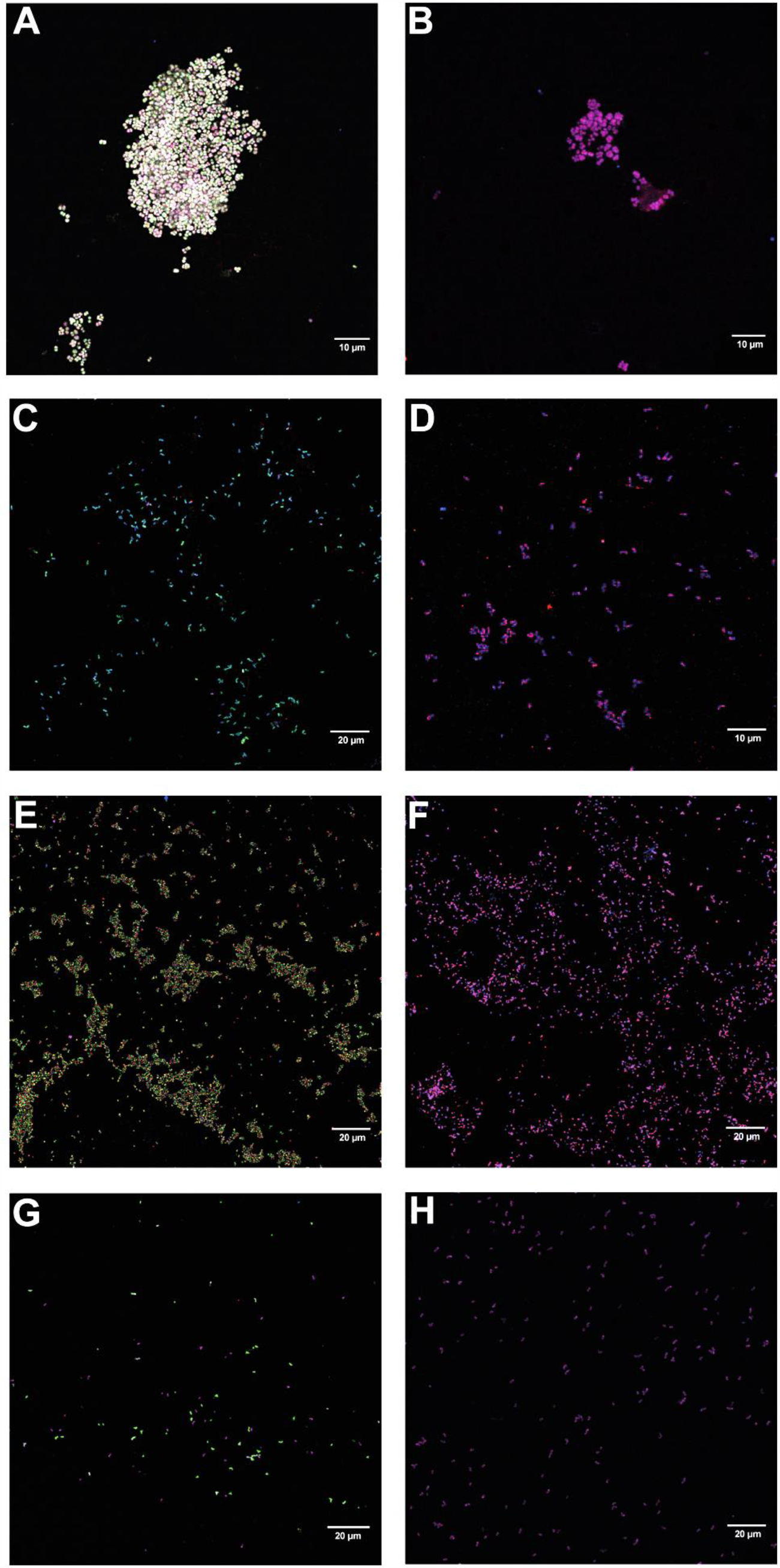
ABPP-based fluorescent labelling of catalytically active methane oxidizing bacteria. (A, C, E, G) 1,7OD pre-incubated and (B, D, F, H) 1,7OD untreated (control) cells of (A, B) *Methylotetracoccus oryzae*, (C, D) *Methylosinus sporium* M29, (E, F) *Methylacidiphilum fumariolicum* SolV, (G, H) *Methylocella tundrae*. Cells are stained with the AMO/MMO-labelling protocol (green) and FISH probes specific for (A, B) Gammaproteobacteria (Gam42a; red), (C, D, G, H) Alphaproteobacteria (Alf0001b, Alf0968; red), (E, F) Verrucomicrobia (EUB338 III; red), and (A-H) all bacteria (EUB338mix).

The efficiency and specificity of the ABPP-based AMO labelling was verified using pure cultures of the following nitrifying microorganisms: (i) *Nitrosomonas europaea*, a well-studied canonical ammonia-oxidizing bacterium, (ii) *Nitrospira inopinata*, the only comammox bacterium available as pure culture (58), (iii) *Nitrospira moscoviensis*, a canonical nitrite-oxidizing bacterium and (iv) *Nitrosocosmicus franklandus*, an ammonia-oxidizing archaeon. Efficient staining of the *N. europaea* and *N. inopinata* cells was observed when 1,7OD-treated cells were subjected to the CuAAC reaction (Figure 2A, C). In contrast, no fluorescent signal was obtained when the same cells were not treated with 1,7OD (Figure 2B, D), or in the canonical, nitrite-oxidizing *N. moscoviensis* (Figure 2F). However, also *N. franklandus* cells did not show any fluorescent labelling (Figure 2E), indicating that the ABPP-based protocol was not able to label the archaeal AMO enzyme.

Although it has been reported that MMOs and AMOs exhibit similar inhibition patterns (35) and it was shown that both sMMO and pMMO enzymes can also be inactivated by acetylene as well as other alkyne compounds (47, 51, 52). The effect of 1,7OD on methane-oxidizing activity has not been studied yet. To test whether the 1,7OD-based ABPP protocol could also label MMOs the following pure cultures were subjected to our labelling protocol: (i) *Methylotetracoccus oryzae*, a type Ib methanotrophic gammaproteobacterium (56); (ii) *Methylosinus sporium*, a type II methanotrophic alphaproteobacterium (54); (iii) *Methylacidiphilum fumariolicum*, an acidophilic methane oxidizer of the phylum *Verrucomicrobia* (6); and (iv) *Methylocella tundrae*, an alphaproteobacterial methanotroph encoding only sMMO (55). Specific 1,7OD-dependent fluorescent staining of all tested methanotrophic cultures indicated efficient ABPP-based labelling of MOB, regardless of enzyme type and phylogenetic affiliation (Figure 3).

### Correlation of the AMO/MMO staining with the bacterial growth stage

Laboratory cultivation conditions poorly reflect the conditions microorganisms encounter in natural and engineered ecosystems. In these environments, fluctuations in substrate and nutrient availability rarely permit the continuous growth of microorganisms, including AOB and MOB (95). To investigate the potential effect of the bacterial growth stage on the efficiency of the AMO-based staining, a *N. europaea* pure culture was cultivated in the presence of 10 mM ammonium and ammonia-oxidizing activity and growth were monitored over 21 days. The culture stoichiometrically oxidized ammonium to nitrite (Figure 4A) and biomass samples taken at different time points were subjected to the ABPP-based protocol. The AMO-derived fluorescent signal intensity reached its maximum at the start of the exponential growth phase (day 7) and decreased when the culture entered early stationary phase, but maintained a low signal intensity (35% compared to the maximum intensity) even in late stationary phase (Figure 4B). Throughout all growth stages of the culture a high percentage (>90%) of cells was stained, with a small decrease (16%) in staining efficiency observed only in late stationary phase (Figure 4B).

**Figure 4.**
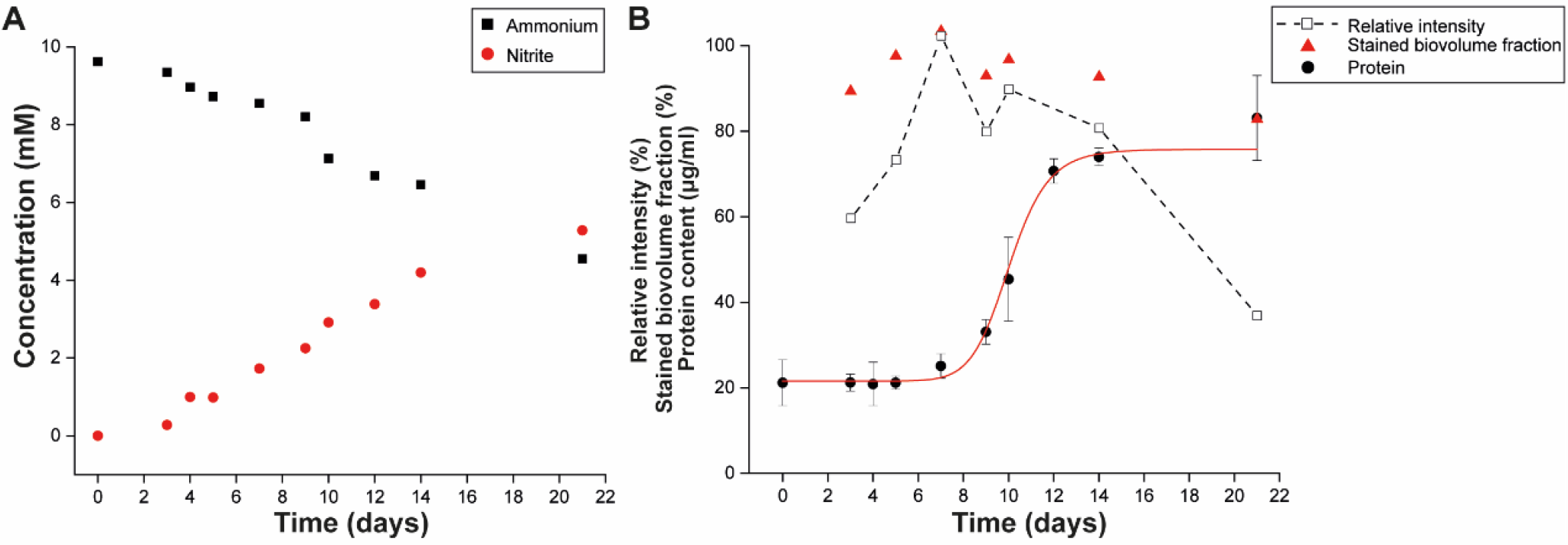
Correlation of the AMO/MMO-derived fluorescent signal intensity and staining efficiency with the growth stage of *Nitrosomonas europaea*. (A) Ammonia consumption and nitrite production activity observed in the culture during the incubation period. (B) Growth of the bacterial culture as indicated by protein content and its relation to the relative intensity of the fluorescent signal observed, as well as the biovolume fraction of the culture stained by the AMO/MMO-based protocol. Bacterial growth data are fitted to a Hill equation. Error bars represent standard deviation of technical triplicates of protein measurements.

### Subcellular localization of active AMO and MMO enzymes

AMO and pMMO are known to be membrane-integral enzymes, with their active center on the periplasmic face of the cytoplasmic membrane or the intracytoplasmic membrane (ICM) systems (96, 97). In contrast, the sMMO is a soluble cytoplasmic enzyme (37). To confirm these subcellular localizations, pure cultures of the canonical ammonia oxidizer *N. europaea*, the comammox bacterium *N. inopinata*, the type Ib methanotroph *M. oryzae* (containing only pMMO), and the methanotroph *M. tundrae* (containing only sMMO) were subjected to the ABPP-based AMO/MMO-staining protocol in combination with FISH and subsequent deconvolution fluorescence microscopy (Figure 5). In *N. europaea* the AMO label was observed mainly in a thick layer surrounding the cytoplasm (Figure 5A), coinciding with the localization of the cytoplasmic membrane and the peripheral organization of the ICM system in *Nitrosomonas* (97-99). In *M. oryzae* the ICMs are arranged as stacks of vesicular discs located in the central cytoplasm (96), which agrees with the observed strong pMMO fluorescent staining on apparently organized structures within the cytoplasm (Figure 5B). As expected, in *M. tundrae* the sMMO labelling overlapped with the cytoplasmic 16S rRNA-derived signal (Figure 5C). Unfortunately, the small cell size (≤ 1 μm) of *N. inopinata* precluded an unambiguous localization of the AMO-derived fluorescent signal (data not shown).

**Figure 5.**
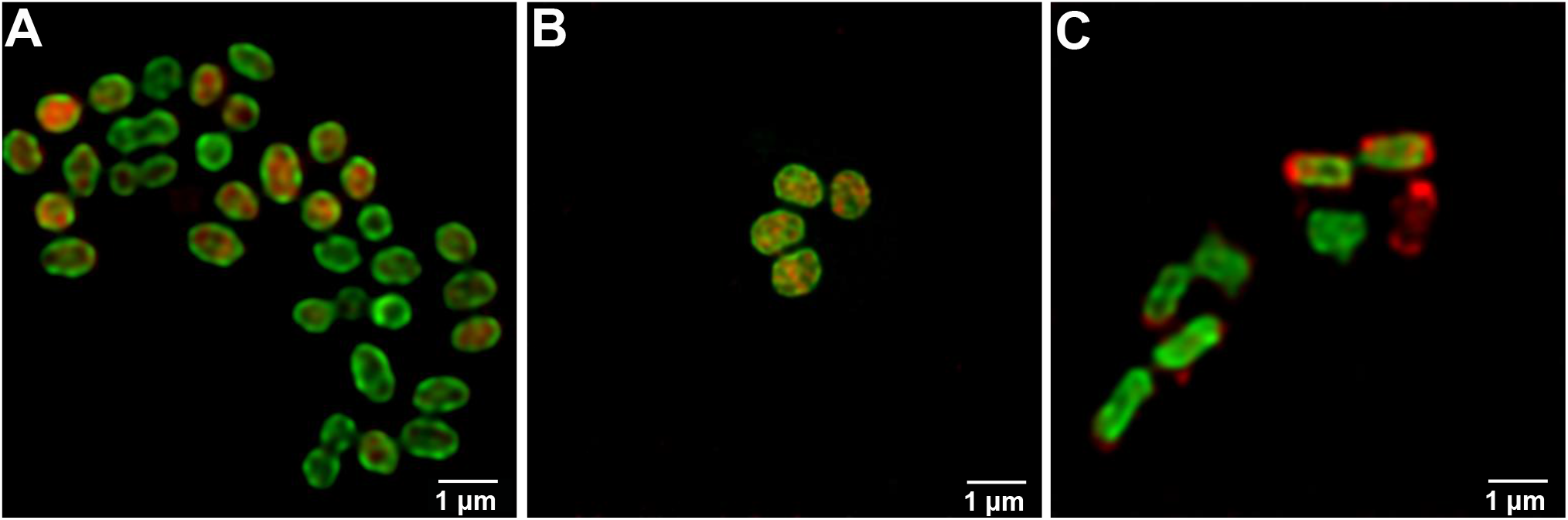
Subcellular localization of the AMO/MMO-derived fluorescent signal using deconvolution confocal microscopy. Cells of (A) *Nitrosomonas europaea*, (B) *Methylotetracoccus oryzae* and (C) *Methylocella tundrae*, are stained with the fluorescent AMO/MMO protocol (green) and specific (A) AOB (Nso190, Neuo653, Nso1225; red), (B) gammaproteobacterial (Gam42a; red) or (C) alphaproteobacterial (Alf0001b, Alf0968; red) FISH probes.

In order to examine the localization of active AMO and pMMO enzymes in more detail, cells of *N. europaea* and *M. oryzae* were subjected to a modified ABPP-based AMO/MMO-staining protocol. In this experiment, the CuAAC reaction was used to biotinylate the AMO/pMMO enzymes within active bacterial cells, with subsequent immunogold labelling and transmission electron microscopy, thus providing high-resolution information on the subcellular localization of the complexes (Figure 6). In *N. europaea*, 74% of the immunogold labelling indicated a membrane localization of the AMO complex, and the majority of these gold particles (49%) appeared to face the periplasmic space, with only 13% oriented towards the cytoplasm (Figure 6B-C, Table S3). In accordance to the proposed localization of the alkynylation site in the MOB *Methylococcus capsulatus* (47), 61% of gold labelling *M. oryzae* depicted a membrane localization, with 71% of these membrane-associated labels on the periplasmic side of the membrane (Figure 6E-F, Table S3). In both cases gold particles were seldomly found in the cytoplasm or outside the cells, indicating the high specificity of the labelling protocol (Figure 6, Table S3).

**Figure 6.**
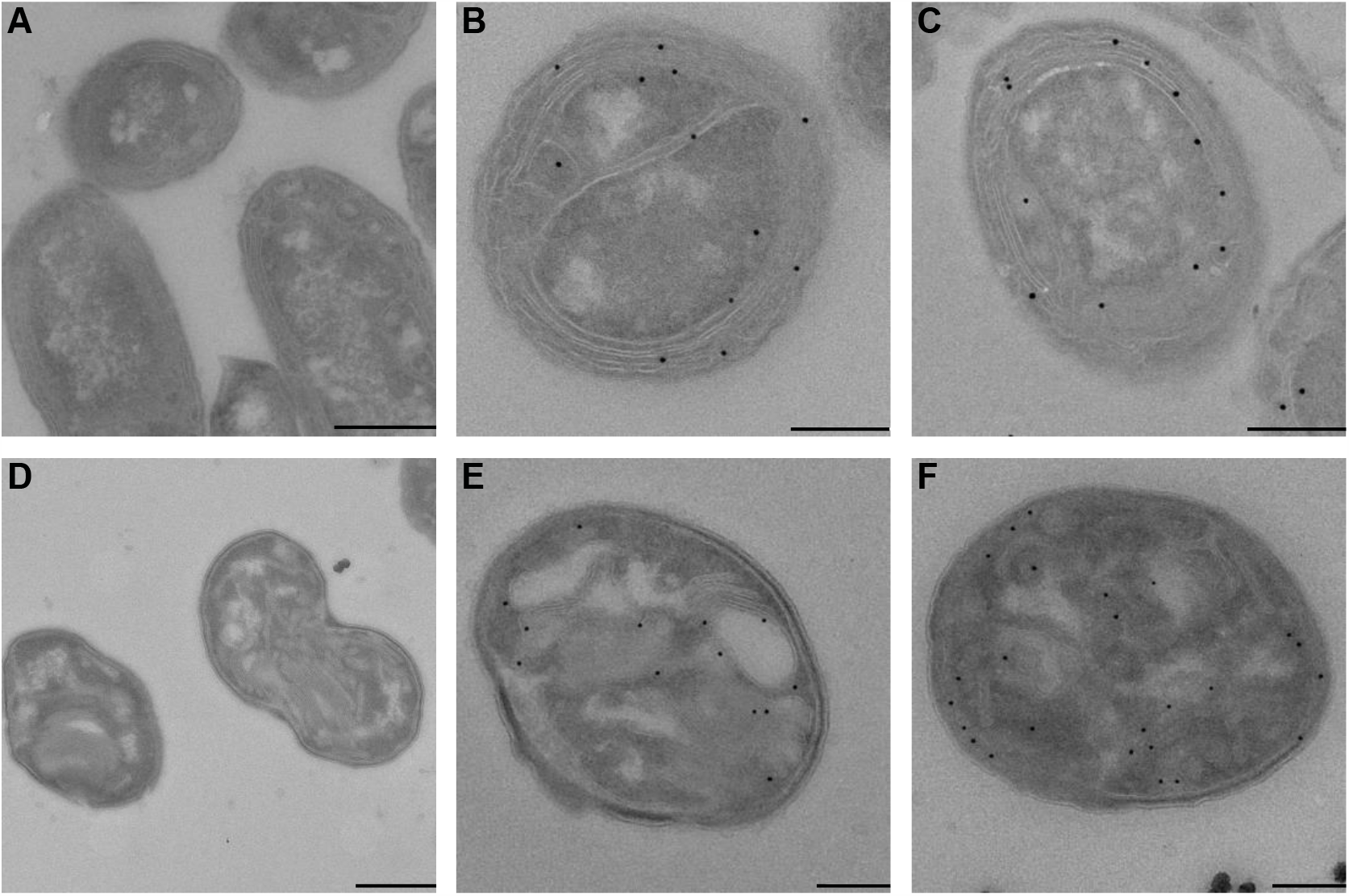
Immuno-gold labelling of active AMO/MMO enzymes using transmission electron microscopy. (A, D) 1,7OD untreated (control) and (B, C, E, F) 1,7OD pre-incubated cells of (A-C) *Nitrosomonas europaea* and (D-F) *Methylotetracoccus oryzae*. Scale bars correspond to 0.5 μm.

### In situ ABPP-based fluorescent labelling of ammonia oxidizers in complex communities

The ability of the ABPP-based AMO/MMO staining protocol to label ammonia oxidizers present in complex microbial communities was assessed in nitrifying enrichment cultures and activated sludge samples. First, we tested our newly developed protocol on an enrichment culture of *Ca*. N. nitrosa and *Ca*. N. nitrificans (2) and a nitrifying co-culture containing canonical AOB, comammox and canonical *Nitrospira*. Furthermore, we demonstrated that combination of the ABPP-based protocol with 16S rRNA-targeted FISH allowed the simultaneous phylogenetic identification of the ammonia-oxidizing bacterial populations in all samples (Figure 7).

**Figure 7.**
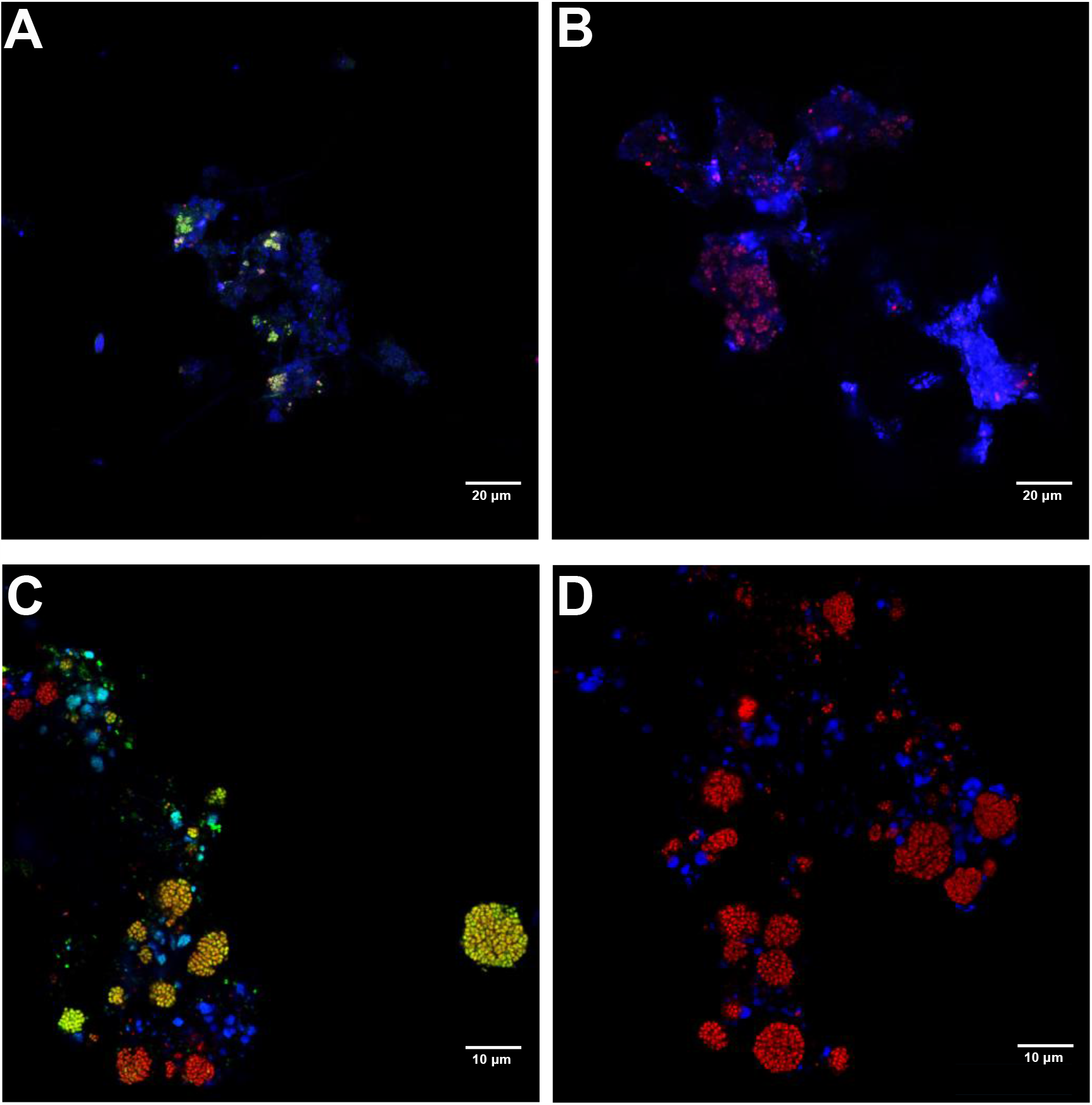
ABPP-based fluorescent labelling of catalytically active ammonia-oxidizing bacteria present in complex microbial communities. (A, C) 1,7OD pre-incubated and (B, D) 1,7OD untreated (control) cells of (A, B) a *Ca*. N. nitrosa and *Ca*. N. nitrificans and (C, D) a nitrifying enrichment culture. Cells are stained with the AMO/MMO-labelling protocol (green signal) and FISH probes specific for *Nitrospira* (Ntspa662, Ntspa712; shown in red [A, B] and in blue [C, D]), (C, D) AOB (Nso190, Neu653, Nso1225; in red) and (A, B) all bacteria (EUB338mix; in blue).

The ABPP-based protocol was able to specifically and efficiently stain both types of ammonia-oxidizing bacteria present in these mixed communities. AMO-derived fluorescent labelling of comammox bacteria was observed both in the enrichment culture containing *Ca*. N. nitrosa and *Ca*. N. nitrificans (Figure 7A, B), and in the nitrifying enrichment (Figure 7C, D). In the latter, betaproteobacterial canonical AOB were also efficiently double-stained by the ABPP-based protocol and FISH, while some *Nitrospira* were detected by FISH only, presumably corresponding to canonical *Nitrospira* present in this sample.

### Targeted metagenomics

In combination with fluorescence-activated cell sorting, our ABPP-based staining protocol could be employed for targeted metagenomics of AMO and MMO-containing bacteria present in complex environmental samples. To prove the feasibility of this approach, we tested the protocol on two samples of different complexity, a nitrifying enrichment culture and activated sludge from a full-scale wastewater treatment plant.

Successive to the application of the ABPP protocol and floc disruption, biomass of the nitrifying enrichment cultures was used for fluorescence-activated cell sorting, with pooled collection of all positive sorting events (Figure S1). This fraction was subjected to DNA extraction and metagenomic sequencing, along with an untreated biomass sample and a range of controls to test for potential biases in the protocol (Table S2). Subsequent metagenomic consensus binning allowed the recovery of three high quality (>92% completeness and <3.7% redundancy) metagenome assembled genomes (MAGs) of ammonia-oxidizing microorganisms from the nitrifying enrichment, corresponding to one *Nitrosomonas* (maxbin2.003) and two *Nitrospira* species (metabat2.63_sub, metabat2.21) that were strongly enriched in the metagenome of the sorted sample (Figure S2, Table S4). Of the two recovered *Nitrospira* MAGs, only metabat2.21 contained an *amoA* sequence, whereas this gene was absent in MAG metabat2.63_sub. However, an unbinned contig containing an *amoA* sequence showed similar coverage to this MAG across all samples, suggesting its association with this *Nitrospira* MAG. All MAGs corresponding to non-ammonia-oxidizing microorganisms were de-enriched in the targeted metagenome (Figure S2, Table S4), indicating a successful targeted retrieval of AMO-containing bacteria.

In order to demonstrate the efficacy of the labelling and sorting protocol in a manner unbiased by metagenomic binning approaches and different sequencing depths, we used metagenomic read mapping to assembled and extracted *amoA* as functional marker for ammonia-oxidizing microorganisms, and *rpoB*, a conserved single-copy phylogenetic marker gene to detect and differentiate all members of the microbial community. In total, ten genes belonging to the copper-containing membrane-bound monooxygenase family (Pfam PF02461) were identified in the co-assembled contigs retrieved from the nitrifying enrichment: three were highly similar to *amoA* of *Nitrosomonas* and two of comammox *Nitrospira* (>96% amino acid identity), while the remaining five were most similar to unresolved genes within proteobacterial lineages not known to oxidize ammonia. These latter genes belonged to a group of putative hydrocarbon monooxygenases (HMO) (100, 101) and became undetectable in the sorted sample, and thus were excluded from further analysis. A total of 48 *rpoB* genes were recovered and analyzed, including two affiliated with *Nitrosomonas* (>90% amino acid identity), two with comammox *Nitrospira* (>97% amino acid identity), and six with canonical *Nitrospira* (>97% amino acid identity). For both marker genes belonging to the dominant ammonia oxidizers in the bioreactor, the normalized read abundances increased in the sorted samples compared to the unsorted samples – *Nitrosomonas* and comammox *Nitrospira amoA* increased 41- and 13-fold, and their *rpoB* 86- and 12-fold, respectively (Figure 8). This increase in abundance was decoupled from the total number of reads per sample recruited to the co-assembly, indicating the changes were not due to variations in sequencing depth (Figure 8). Finally, the abundance of non-ammonia-oxidizing community members’ *rpoB* genes decreased from 93.6% of the total normalized read abundance to 12.5%, indicating a strong de-enrichment of the microbial community members lacking AMO (Figures 8 and S2). Therefore, the ABPP-based fluorescent labeling coupled to cell sorting successfully and specifically enriched the ammonia-oxidizing microorganisms present in the nitrifying bioreactor.

**Figure 8.**
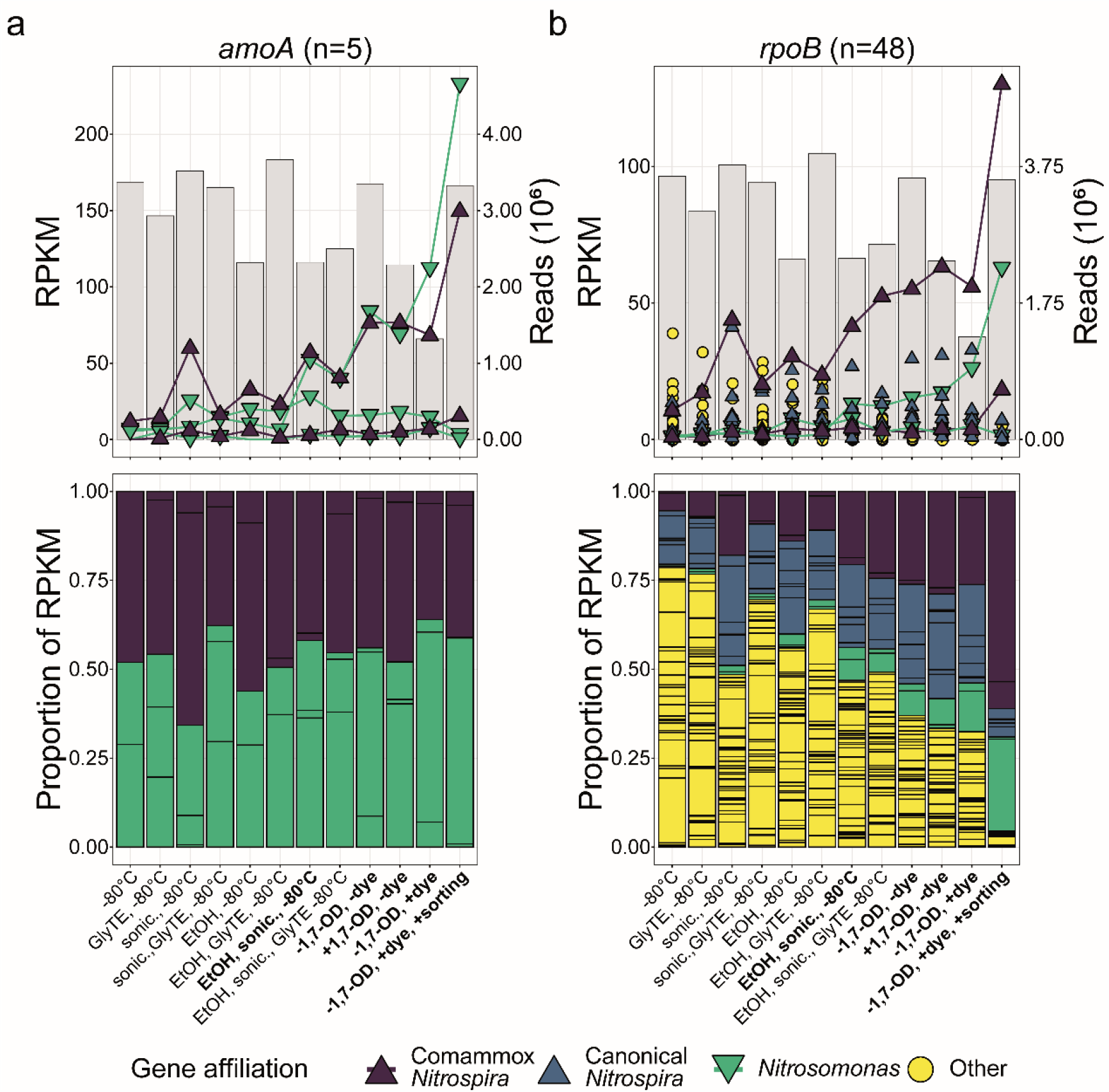
Normalized and relative abundances of (A) *amoA* and (B) *rpoB* genes in metagenomes of untreated, controls, and sorted samples. The top row shows the normalized read abundances in units of Reads Per Kilobase gene length per Million mapped reads (RPKM) as symbols colored by taxonomic affiliation, and the total number of reads mapped to the co-assembly as gray bars. The same gene for the taxa of interest, *Nitrosomonas* and comammox *Nitrospira*, are emphasized and connected with lines. The bottom shows the same RPKM data as the top row as a proportion of the total RPKM for the respective gene in the metagenomic sample, with each individual colored box representing the relative abundance of a single gene from the co-assembly. The final treatment preceding the CuAAC reaction is indicated in bold along with the controls for the inactivation and dyeing, and the final sorted sample.

However, the ABPP-based labelling and sorting approach did reveal biases and limitations. First, there appeared to be enrichment of nitrifying bacteria due to treatment steps prior to sorting, particularly among the sonicated and ethanol-treated samples, but also by the CuACC reaction itself (Figure 8). These effects may be due to their tendency to grow as microcolonies within biomass aggregates, which may protect them from the effects of sonication, ethanol fixation, and the CuAAC reaction (92-94). Second, the apparent most abundant ammonia-oxidizer after ABPP-based labelling and cell sorting depended on the marker gene, as a *Nitrosomonas*-like canonical AOB appeared to be most highly enriched based on *amoA* abundance increases, while this was a comammox *Nitrospira* in the *rpoB* analysis. The slopes of linear correlations (r^2^ >0.95, Figure S3) of these most abundant *amoA* and *rpoB* genes support genomic evidence that this discrepancy may be an effect of the copy number of *amoA*, i.e. *Nitrosomonas* typically encode three copies of *amoA* but comammox *Nitrospira* only one (102). Third, the larger increases in abundance of both *Nitrosomonas* marker genes compared to comammox *Nitrospira* suggest that the technique may slightly favor *Nitrosomonas*-like AOB. Fourth, some *amoA* and *rpoB* affiliated with AOB were either poorly enriched or even de-enriched, which makes it tempting to speculate on lifestyles independent of ammonia oxidation that might result in absence of the AMO enzyme in these cells (Figure 8). While important to acknowledge and account for these caveats, metagenomic recovery and abundance estimations clearly showed effective enrichment of ammonia-oxidizing microorganisms by the ABPP-labelling protocol coupled to fluorescence-activated cell sorting.

When applied to activated sludge sampled at a municipal wwtp, the ABPP-based protocol in combination with cell sorting (Figure S1) and metagenomic sequencing facilitated a 128-fold increase in coverage of the most abundant *Nitrosomonas* species MAG (metabat2.58) in comparison to the untreated sample, demonstrating its successful and targeted enrichment (Figure 9, Table S5). In general, canonical ammonia oxidizers were of extremely low abundance in this wwtp, and this MAG only contributed 0.14% of all mapped reads in the metagenome of the untreated sample. After cell sorting and sequencing, it was enriched to account for 8.64% of the mapped reads, which allowed the recovery of a high-quality draft genome with estimate completeness of 100% and 0.81% redundancy (Figure 9, Table S5). A second MAG (metabat2.38) affiliated with the genus *Nitrosomonas* accounted for 0.12% and 0.96% of mapped all reads in the samples before and after sorting respectively, corresponding to an 8-fold increase in coverage in the sorted sample (Table S5). This clearly demonstrates the efficacy of our ABPP-based protocol for the targeted recovery of ammonia-oxidizing bacteria, even when they constitute a minute fraction of the original community.

**Figure 9.**
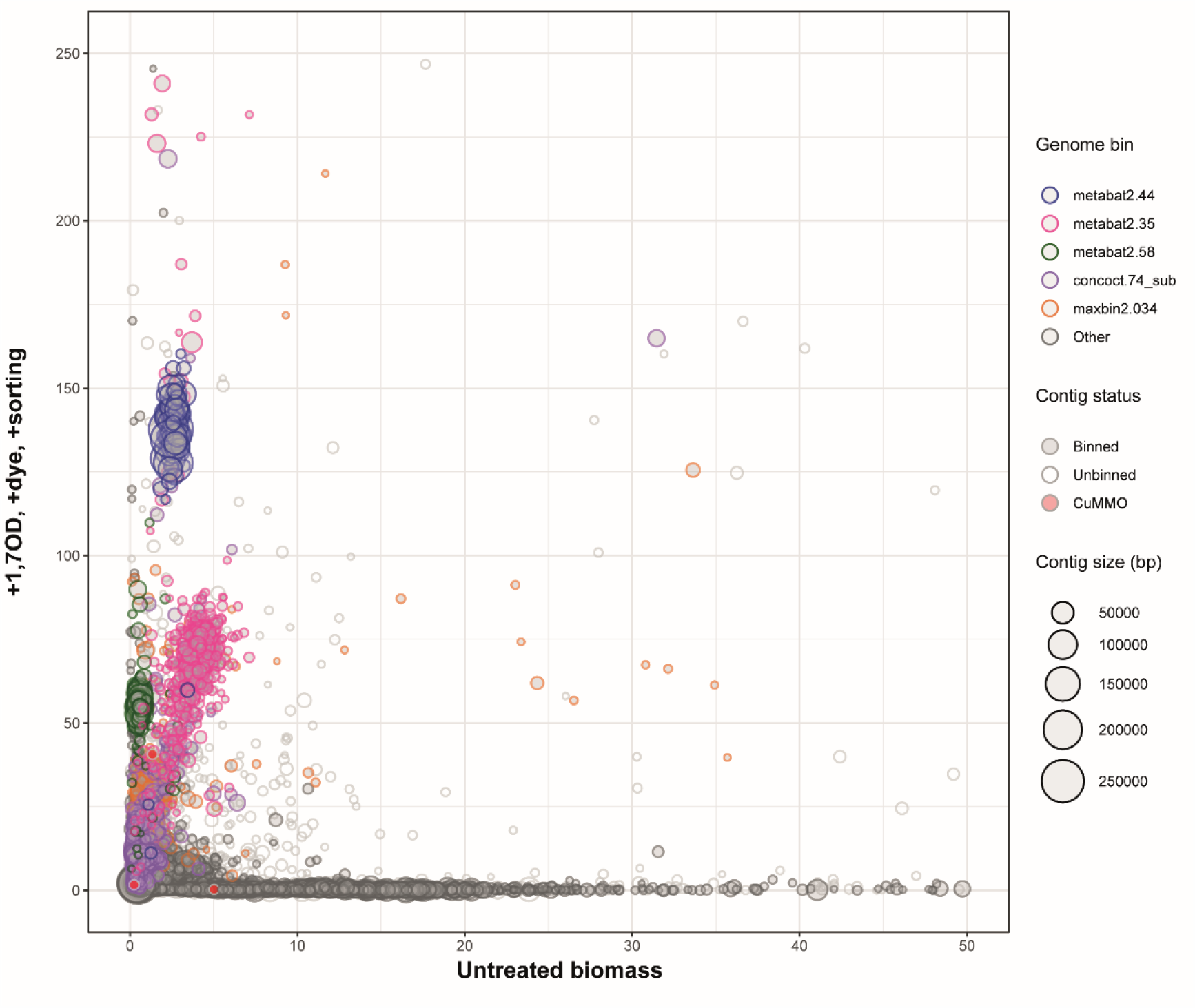
Differential coverage plot showing the abundance of contigs in the untreated activated sludge sample in comparison to the ABPP labelled and sorted sample. Each circle represents a metagenomic contig, the circle size corresponds to its length. The five most abundant MAGs in the ABPP-positive and sorted sample are indicated by color, while the red filling indicates contigs on which genes encoding CuMMO (AMO or MMO) subunits have been detected.

Surprisingly, a high percentage (65.2%) of the total reads obtained for the sorted activated sludge sample were affiliated with putative glycogen-accumulating organisms (GAOs) of the family *Competibacteraceae* (Tables S5). We obtained in total 7 MAGs (>70% completeness) affiliated with this family, which showed an increase in coverage from 8.9 to 52-fold in the sorted compared to the untreated metagenomic sample (Figure 9, Table S5). However, none of these MAGs contained genes encoding AMO or MMO subunits, although some unbinned *amoA* sequences were present in the full assembly. Thus, we performed manual binning and bin refinement using a new assembly of the sequencing reads of the sorted sample only, but using differential coverage information from all samples. These refined MAGs were then screened for genes involved in ammonia or methane oxidation, or genes which otherwise might explain the enrichment of these GAOs by the ABPP-based sorting method. The manual binning allowed us to link the observed *amo* genes to the dominant *Nitrosomonas*-affiliated MAG, but revealed no traces of an ammonia-oxidizing potential for any of the *Competibacteraceae* family members (three *Contendobacter* and one *Competibacter* species with an estimated genome completeness >70%; Table S6), and also MMO genes were not detected in these MAGs. This indicates that the observed enrichment was not a result of an AMO/MMO-based labelling and the only enzymes encoded by these *Competibacteraceae* MAGs known to interact with alkynes and thus potentially explaining their fluorescent labelling (Figure S4) were nitrogenases (103).

Consequently, to test for potential labelling of the nitrogenase complex, we incubated pure cultures of the diazotroph *Krypidia spormannii* FAVT5 (59) and the methanotroph *M. oryzae* (56) under nitrogen-fixing conditions. Subsequently, both cultures were subjected to the ABPP-based protocol. As *M. oryzae* contains a pMMO, staining of these cells was observed, but compared to a culture grown in the presence of nitrate as N-source in the medium, the staining intensity was even slightly reduced (Figure S5). For *K. spormannii* FAVT5 the amount of signal observed after the CuAAC reaction was just above background, corresponding to approximately 11% and 25% of the signal obtained from active *N. europaea* and *M. tundrae* cultures, respectively (Figure S5). Together, these results indicate that the presence of an active nitrogenase does not contribute to the ABPP-based labelling, making the reason for the observed staining of the *Competibacteraceae*-affiliated MAGs unclear.

## Discussion

ABPP-based protocols have been successfully employed for the study of many microbial protein families, such as proteases, kinases, hydrolases and glycosidases (8, 17). However to date, they have not been applied for the labelling of whole cells within complex microbial communities, mainly due to the presence of the microorganisms within a complex undefined matrix. This poses major challenges for employing the CuAAC reaction, probe penetration into the cells, and labelling specificity and efficiency (104). The ABPP-based protocol developed in this study, to our knowledge for the first time, overcomes these challenges and allows the fluorescent visualization of catalytically active enzymes *in situ*. The protocol makes use of the alkyne 1,7OD, which serves as a bifunctional enzyme probe for the specific detection of AMO and MMO enzymes. The feasibility of this approach was previously demonstrated *in vitro* for the AMO of canonical AOB (19), but had not been tested for labelling whole cells, phylogenetically distinct ammonia oxidizing microorganisms, or MMOs of methanotrophs. Generally, *n*-terminal and sub-terminal alkynes are known mechanism-based inactivators of both AMO (48, 49) and MMO enzymes (46, 51). Here, we could show that 1,7OD inhibits ammonia oxidation in comammox *Nitrospira* (Figure 1) as efficiently as was reported for canonical AOB (19). Furthermore, strong labelling of different MOB harboring either the soluble or particulate MMO (Figure 3) indicated that also methanotrophs will be inhibited by this diyne. Application of the *in situ* ABPP-based protocol on pure and enrichment cultures of different canonical AOB, comammox *Nitrospira*, and MOB resulted in their efficient and specific labelling (Figures 2 and 3), which directly conveyed information on the functional potential of the detected microorganisms in contrast to conventional *in situ* detection methods like FISH. Using high-resolution deconvolution light microscopy as well as electron microscopy, the ABPP-based protocol furthermore permitted visualization of the subcellular localization of the AMO and MMO enzymes (Figures 5 and 6). In accordance with previous studies (55, 98, 99), the AMO and pMMO-derived fluorescent signals were mostly localized along the cytoplasmic membrane and on the intracytoplasmic membrane stacks in *N. europaea* and *M. oryzae*, respectively, whereas a cytoplasmic localization of the sMMO-derived signal was observed for *M. tundrae*. Furthermore, the immuno-gold labelling data obtained in this study suggested that the alkynylation sites for both the AMO and pMMO complexes were found on the periplasmic face of the ICMs. This congruence with the expected enzyme localizations further verified the suitability of the ABPP-based protocol for the specific labelling of AMO/MMO enzymes, and opens up possibilities for studying differences in enzyme distribution along intracellular structures in AOB or MOB types with divergent cell morphologies. Simultaneously, the biotinylation of these enzymes via the ABPP-based protocol as tested here provides a new methodology for the profiling, affinity purification and study of AMO/MMO enzymes.

In most natural and engineered systems microorganisms rarely encounter optimal growth conditions and might be in very different growth stages. Previous studies have identified preservation of basal activity levels of ammonia and methane oxidizers as a general adaptation mechanism used to cope with alternating conditions and fluctuating substrate supply (95, 105). This indicates that they maintain a basal AMO/MMO content throughout their life cycle. Quantification of ABPP-derived fluorescent signals in a *N. europaea* pure culture showed a strong correlation between signal intensity and growth state (Figure 4). As expected, the strongest signals were observed during the exponential growth phase, corresponding to a high AMO content of the cells. However, detectable ABPP-based labelling was observed even in the late stationary phase of the culture, demonstrating that basal AMO contents are maintained also in non-replicating cells, which however did not encounter substrate limitation. Thus, our ABPP-based protocol apparently is suitable for the detection of ammonia and methane-oxidizing bacteria also under suboptimal growth conditions.

As the ABPP-based protocol can be combined with FISH, it allows to directly link the functional potential of a microorganism to its identity *in situ*, which potentially even can lead to the identification of novel ammonia or methane oxidizers. The need for simple detection methods not requiring *a priori* knowledge on the identity of microorganism was showcased by the recent discovery of complete nitrification within members of the genus *Nitrospira* (1, 2). While *amoA*-targeting PCR-based methods have successfully been employed to detect the presence and abundance of comammox bacteria in complex microbial communities or environmental samples (106), the distinction between canonical and comammox *Nitrospira* based on their phylogenetic affiliation is still problematic. Consequently, it also is not possible to reliably detect comammox *Nitrospira* in 16S rRNA-targeted FISH, which hampers research on their role in full-scale biotechnological systems like drinking and wastewater treatment plants. In combination with FISH, the ABPP-based AMO/MMO staining method described here overcomes this limitation and enables an easy and reliable differentiation of ammonia and nitrite-oxidizing *Nitrospira* in complex microbial communities (Figure 7).

Unfortunately, it was not possible to efficiently stain the archaeal AMO using 1,7OD as bifunctional enzyme probe (Figure 2E). This is in accordance with the low sensitivity of AOA to longer-chain length (>C_5_) alkynes and their reversible inhibition by 1-octyne (49). Nevertheless, as AOA are sensitive to inactivation by n-alkynes with short carbon backbones (≤C_5_) (19), a modification of the ABPP-based protocol employing a smaller enzyme probe might permit also labelling of AOA. However, the fact that diynes smaller than 1,5-hexadiyne are extremely reactive and difficult to obtain will make the applicability of this method for the detection of AOA challenging.

Generally, most nitrifiers and methane oxidizers are notoriously difficult to cultivate due to their slow growth rates and susceptibility to contamination. Besides classical cultivation attempts, the identification of novel ammonia and methane oxidizers to date was achieved using metagenomic sequencing (1-6, 107-110) and molecular techniques (106, 111, 112). These contemporary cultivation-independent tools have been invaluable for providing insights into the distribution and ecological importance of these functional groups. Despite the fact that metagenomic sequencing has provided a wealth of information on these microorganisms, it is often limited by the ecosystems’ complexity and required resource investments, and can fail to recover sufficient data to reconstruct genomes of low-abundance organisms. Furthermore, the lack of PCR primers able to amplify novel and possibly divergent sequences can hinder the detection of new species with novel metabolic capabilities. Additionally, linking novel sequence types detected in metagenomic or PCR-derived datasets to a specific phylotype often is a highly challenging and time-consuming task. Thus, there is a pressing need for simple detection methods that do not requiring *a priori* knowledge on the identity of microorganism.

To overcome these limitations, we combined the newly developed ABPP-based AMO/MMO staining protocol with cell sorting and subsequent metagenomic sequencing of the sorted subpopulation. Application of this protocol on a nitrifying bioreactor enrichment culture substantially and specifically enriched the multiple types of AOP present in the biomass (Figure 8). While the physical and chemical treatments of the biomass during the ABPP-based protocol were observed to introduce biases resulting in a slight unspecific enrichment of microorganisms like *Nitrosomonas* and *Nitrospira*, which possibly can be explained by an increased cell stability due to their tendency to form microcolonies (113, 114), only the final sorting step of the labelled bacteria resulted in the high and specific enrichment of these AOP (Figure 8).

When applied on activated sludge from a full-scale municipal wwtp, the ABPP-based labelling protocol with subsequent cell sorting resulted in the targeted recovery of a *Nitrosomonas* MAG (Figure 9). This MAG was of very low abundance in the original sample, but the 128-fold increase in coverage achieved by our staining and sorting approach allowed the reconstruction of a high quality genome, demonstrating the power of this ABPP-based protocol.

Surprisingly, also several MAGs belonging to members of the *Competibacteraceae* family were retrieved from the activated sludge sample. *Contendobacter* and *Competibacter* species are glycogen-accumulating organisms, which frequently are encountered in enhanced biological phosphorus removal systems. In these systems, they compete with phosphate-accumulating organisms for resources under the cyclic anaerobic ‘feast’ – aerobic ‘famine’ regime (115) and thus reduce the phosphate removal efficiency of these wastewater treatment systems. Their main metabolism includes synthesis and storage of glycogen and polyhydroxyalkanoates, but they are also able of fermentation, denitrification and nitrogen fixation (116). However, neither the annotation of the obtained MAGs nor existing literature supports the involvement of these *Competibacteraceae* species in the nitrification process or methanotrophy. The reason of their labelling and sorting (Figures S1 and S4) through the ABPP-based protocol thus is unclear and requires future research.

In conclusion, we present a novel ABPP-based protocol that enables the efficient *in situ* labelling of AMO/MMO-containing bacteria and facilitates the targeted retrieval of enriched metagenomes. Although optimization of the protocol for difficult sample types that exhibit high background fluorescence will be necessary, the developed ABPP-based labelling protocol is a significant milestone for integrating phylogenetic and physiological information that can be adapted and applied to complex microbial communities. Coupling this novel method with more common techniques can help reveal the functional differences in biomass aggregates of phylogenetically indistinguishable bacteria, as well as enable the recovery of new genomic representatives critical for engineered processes that account for only a small portion of the microbial community. Therefore, the future application of this ABPP protocol will contribute greatly to an improved understanding of the microorganisms involved in important carbon and nitrogen cycle processes within a variety of ecosystems.

## Acknowledgements

The authors would like to thank C. Hogendoorn, M. Ghashghavi, M.A.R. Kox, C.J. Sedlacek and L. Lehtovirta-Morley for the generous provision of biomass. and T. van Alen and H. Harhangi for technical assistance. We are grateful to H. Koch, M. Wagner and H. Daims for helpful discussions. D.S. and M.S.M.J. were supported by the European Research Council (ERC Advanced Grant Ecomom 339880), J.F., G.S., L.K., M.A.H.J.v.K and S.L. by the Netherlands Organisation for Scientific Research (NWO; Gravitation Grant SIAM 024.002.002, 016.Veni.192.062 and 016.Vidi.189.050).

## Author Contributions

S.L. conceived the presented research. D.S., M.A.H.J.v.K., M.S.M.J. and S.L. planned research. S.L. and M.A.H.J.v.K supervised the project. D.S., G.S., R.J.M., J.F. and L.K. executed experiments and analyzed data. D.S., M.A.H.J.v.K. and S.L. wrote the paper. All authors discussed results and commented on the manuscript.

