## Supplemental Material for "An activity-based labelling method for the detection of ammonia and methane-oxidizing bacteria"

**Supplemental Figures and Tables:**

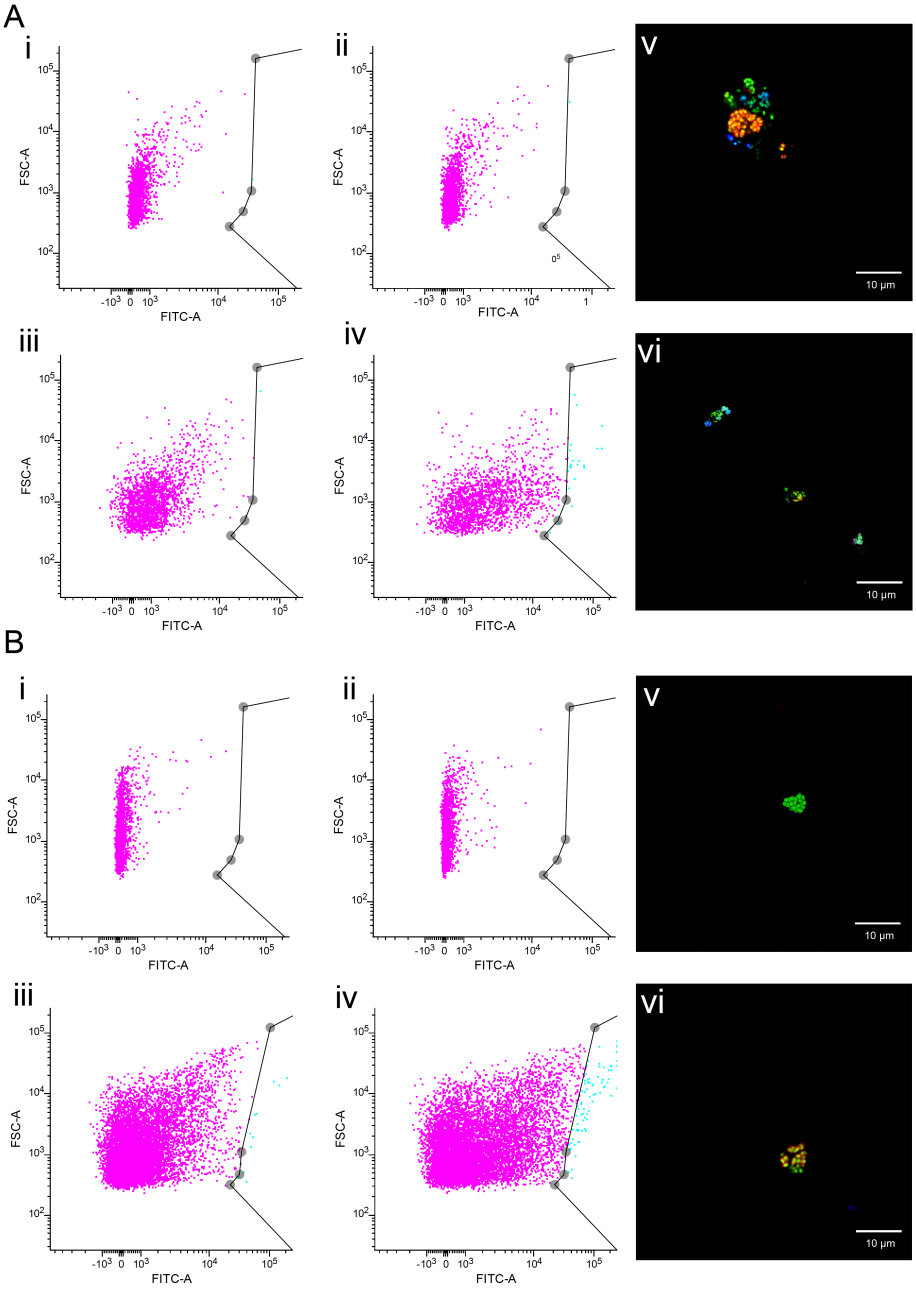


**Figure S1.** Gating of the ABPP-based labelled fluorescent subpopulation of (A) the nitrifying enrichment and (B) activated sludge. (i) biomass incubated without 1,7OD and subjected to a CuAAC reaction without Fluor488-azide, (ii) biomass incubated with 1,7OD and subjected to a CuAAC reaction without Fluor488-azide, (iii) biomass incubated without 1,7OD and subjected to a CuAAC reaction, (iv) biomass incubated with 1,7OD and subjected to a CuAAC reaction, (v, vi) representative FISH pictures [AMO/MMO-belling protocol (green), AOB (Nso190, Neuo653, Nso1225, red) and *Nitrospira* (Ntspa662, Ntspa712, blue)]of the sorted biomass.

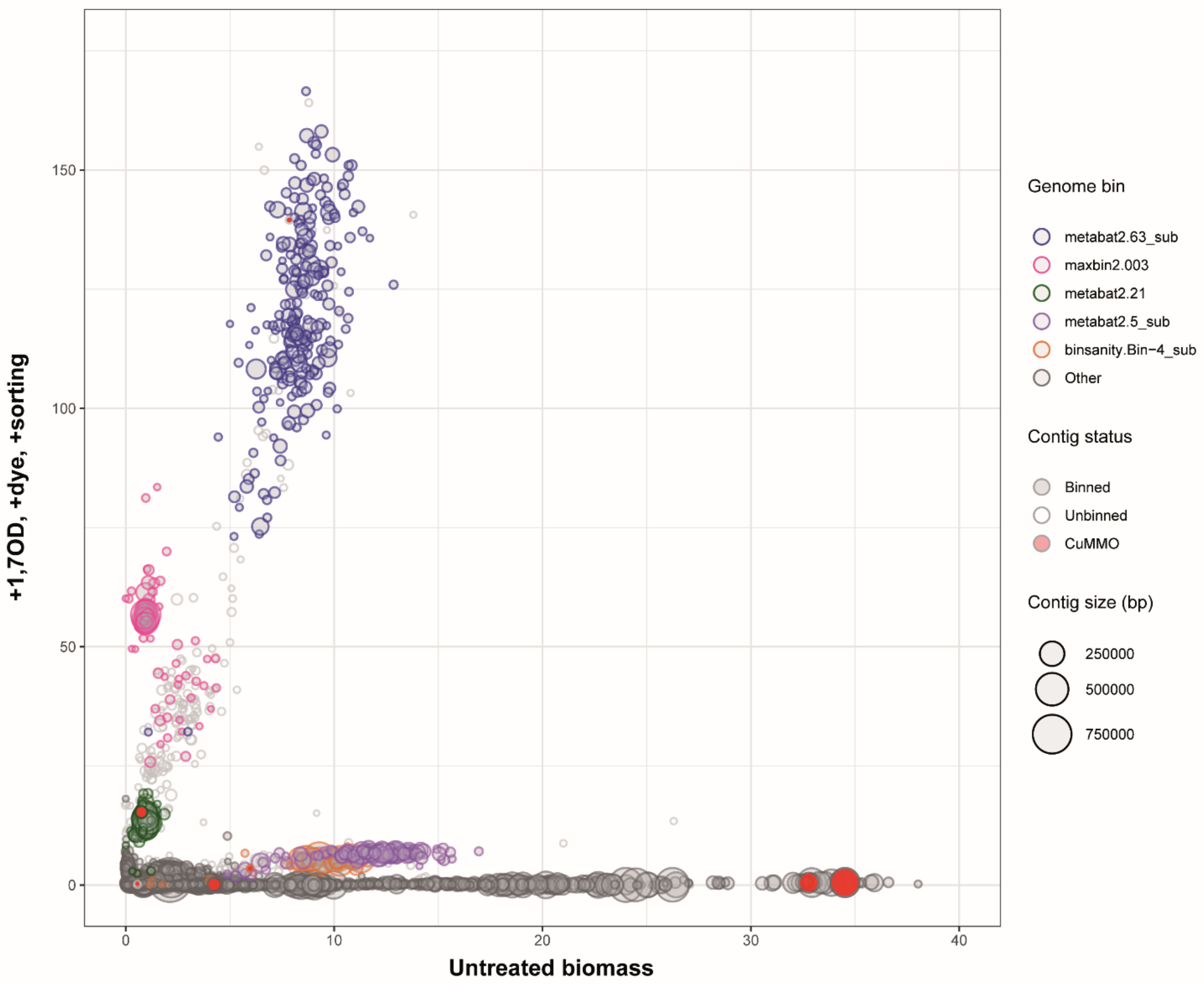


**Figure S2.** Differential coverage plot showing the abundance of contigs in the untreated nitrifying enrichment culture sample in comparison to the ABPP-treated sorted sample. Each circle represents a metagenomic contig, with circle size corresponding to contig length. The five most abundant MAGs in the sorted sample are indicated by color, while the red filling indicates contigs in which copper-containing membrane-bound monooxygenases (AMO, pMMO, HMO) genes have been detected.

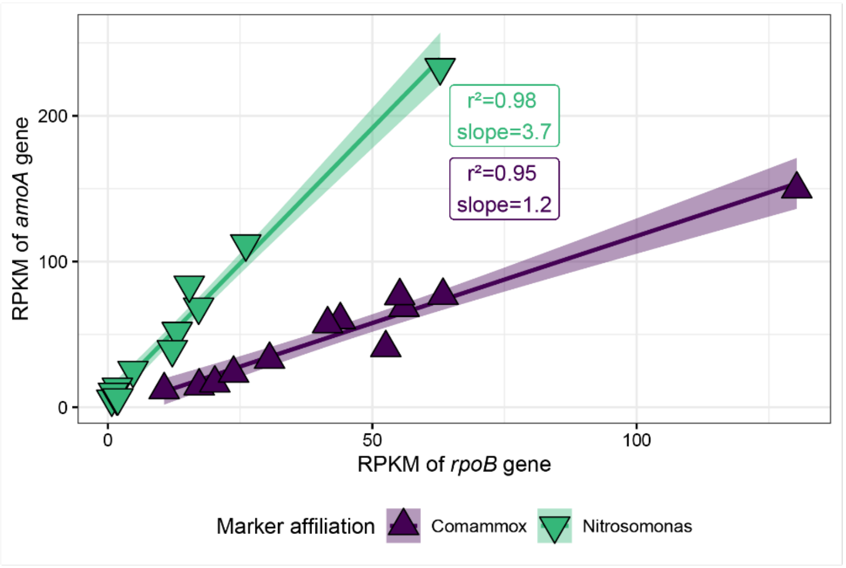


**Figure S3.** Linear correlations of the *amoA* and *rpoB* normalized read abundances of the dominant ammonia-oxidizing bacteria. Each symbol represents the Reads Per Kilobase gene length per Million mapped reads (RPKM) of a gene in one metagenomic sample, colored by taxonomic affiliation to match Figure 9. The linear relationship and the 95% confidence interval are shown as a line and background of matching color, and are labelled with the coefficients and the slope of the line in matching colors.

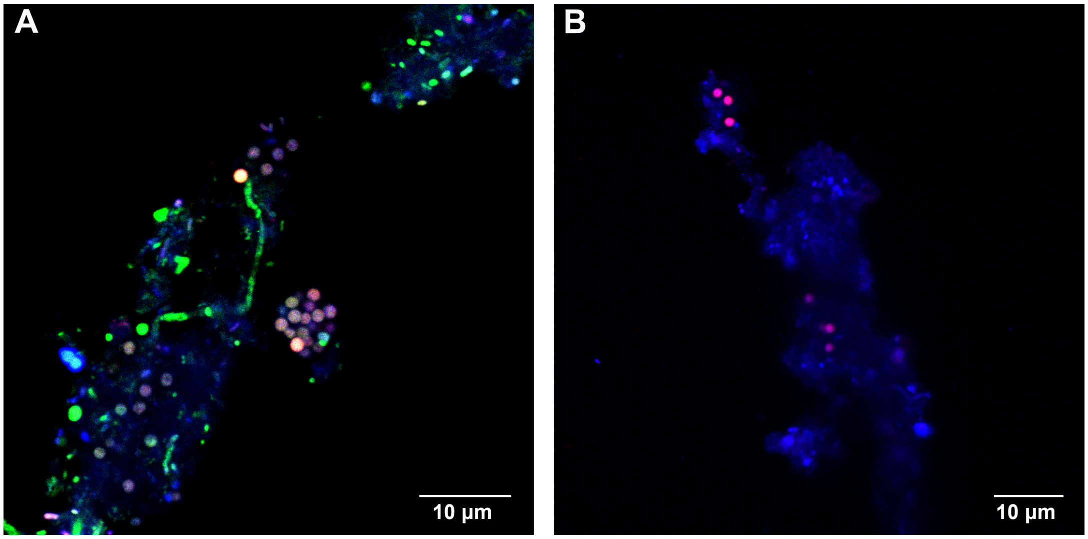


**Figure S4.** ABPP-based fluorescent labelling of (A) 1,7OD pre-incubated and (B) 1,7OD untreated (control) *Competibacteraceae* cells present in activated sludge from a municipal wwtp. Cells were stained with the AMO/MMO-labelling protocol (green) and FISH probes targeting *Competibacteraceae* sp. (GAOQ431, GAOQ989, red) and all bacteria (EUBmix, blue).

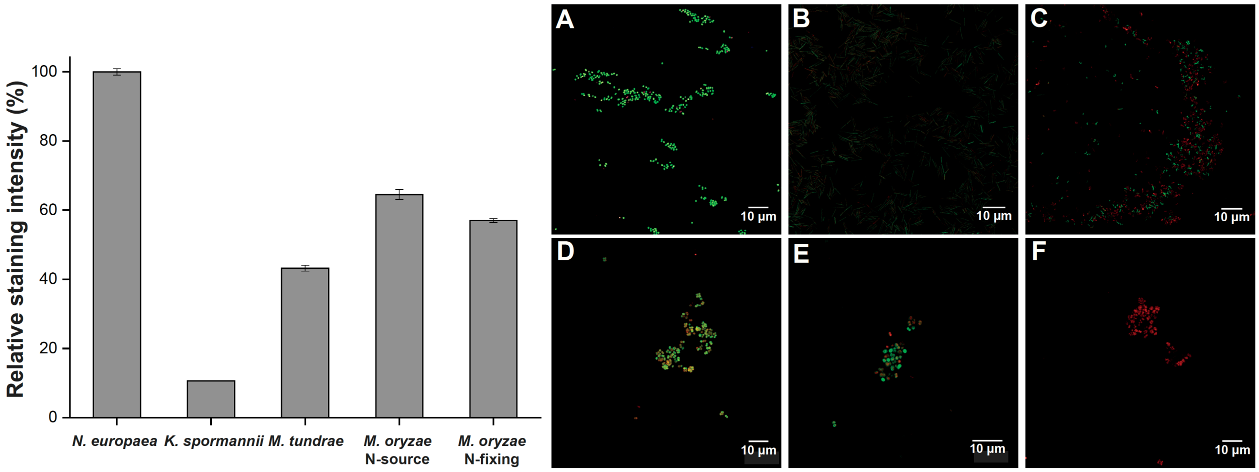


**Figure S5.** Observed relative AMO/MMO-derived fluorescent signal intensity in the canonical AOB *N. europaea*, the nitrogen fixing bacterium *K. spormannii* FAVT5, the sMMO-containing MOB *M. tundrae* and the pMMO-harboring MOB *M. oryzae* grown in the presence of an external nitrogen source or under $N_2$-fixing conditions. Representative pictures of the ABPP-based stained biomass of: (A) *N. europaea*, (B) *K. spormannii* FAVT5, (C) *M. tundrae*, (D, E, F) *M. oryzae* cells grown (D) with external nitrogen source or (E, F) under nitrogen-fixing conditions incubated (E) with or (F) without 1,7OD.

**Table S1.** Specifications of the FISH probes used in this study.

| Probe | Target | % FA | Sequence (5'→3') | Reference |
|---|---|---|---|---|
| **Ntspa662** | *Nitrospira* genus | 35 | GGA ATT CCG CGC TCC TCT | (1) |
| **cNtspa662** | Competitor to Ntspa662 | - | GGA ATT CCG CTC TCC TCT | (1) |
| **Ntspa712** | *Nitrospira* phylum (most members) | 35 | CGC CTT CGC CAC CGG CCT TCC | (1) |
| **cNtspa712** | Competitor to Ntspa712 | - | CGC CTT CGC CAC CGG TGT TCC | (1) |
| **Gam42a** | Gammaproteobacteria | 40 | GCC TTC CCA CAT CGT TT | (2) |
| **Nso190** | Betaproteobacterial AOB | 45 | CGA TCC CCT GCT TTT CTC C | (3) |
| **Nso1225** | Betaproteobacterial AOB | 35 | CGC CAT TGT ATT ACG TGT GA | (4) |
| **Neu653** | *Nitrosomonas* spp. | 35 | GCT GCC ACC CGT AGG TGT | (3) |
| **cNeu653** | Competitor to Neu653 | - | TTC CAT CCC CCT CTG CCG | (3) |
| **EUB338I** | Most Bacteria | 0-80 | GCT GCC TCC CGT AGG AGT | (5) |
| **EUB338II** | *Planctomycetales* | 0-80 | GCA GCC ACC CGT AGG TGT | (6) |
| **EUB338III** | *Verrucomicrobiales* | 0-80 | GCT GCC TCC CGT AGG AGT | (6) |
| **ARCH915 (ARC915)** | Archaea | 20-60 | GTG CTC CCC CGC CAA TTC CT | (5) |
| **GAOQ431** | *Competibacteraceae* | 35 | TCC CCG CCT AAA GGG CTT | (7) |
| **GAOQ989** | *Competibacteraceae* | 35 | TTC CCC GGA TGT CAA GGC | (7) |

**Table S2.** Detailed description of the controls included into the targeted metagenomics approach to determine the potential biases introduced by the different treatment steps of the ABPP-based protocol.

| Treatment / Sample name[a] | Storage at -80°C | GlyTE buffer | Soni-cation | EtOH fixation | 1,7OD addition | CuACC reaction | Azide-Fluor 488 addition | FACS sorting |
|---|---|---|---|---|---|---|---|---|
| **-80°C (untreated)** | ■ | | | | | | | |
| GlyTE, -80°C | ■ | ■ | | | | | | |
| Sonic, -80°C | ■ | | ■ | | | | | |
| Sonic, GlyTE, -80°C | ■ | ■ | ■ | | | | | |
| EtOH, -80°C | ■ | | | ■ | | | | |
| EtOH, GlyTE, -80°C | ■ | ■ | | ■ | | | | |
| EtOH, sonic, -80°C | ■ | | ■ | ■ | | | | |
| EtOH, sonic, GlyTE, -80°C | ■ | ■ | ■ | ■ | | | | |
| **-1,7OD, -dye** | ■ | ■ | ■ | ■ | | ■ | | |
| **+1,7OD, -dye** | ■ | ■ | ■ | ■ | ■ | ■ | | |
| **-1,7OD, +dye** | ■ | ■ | ■ | ■ | | ■ | ■ | |
| **+1,7OD, +dye, sorting** | ■ | ■ | ■ | ■ | ■ | ■ | ■ | ■ |

[a]Treatment controls in bold were included for both environmental samples, all others only for the nitrifying enrichment culture.

**Table S3.** Subcellular localization of the immuno-gold labels in *N. europaea* and *M. oryzae*.

**Table S4.** Metagenomic sequencing information for the nitrifying bioreactor sorting experiment. Details on the taxonomic assignment and quality of the retrieved bins, their non-normalized and sequence-depth-normalized coverage, total number and read percentage aligned to each bin across the different samples are given.

**Table S5.** Metagenomic sequencing information for the wwtp sorting experiment. Details on the taxonomic assignment and quality of the retrieved bins, their non-normalized and sequence-depth-normalized coverage, total number and read percentage aligned to each bin across the different samples are given.

**Table S6.** *De novo* assembled and binning of the data retrieved from the wwtp sorting experiment. Manually refined bins were matched back to the original bins for naming. Details on the taxonomic assignment of each bin, their non-normalized and sequence-depth-normalized coverage, total number and read percentage aligned to each bin across the different samples are given.
